# Hierarchical Gene Cluster Regulation Across Vertebrate Skins: Developmental Control of Keratin Gene Expression

**DOI:** 10.64898/2026.07.14.738566

**Authors:** Wen-Chien Jea, Ping Wu, Chih-Kuan Chen, Cheng-Ming Chuong, Ya-Chen Liang

## Abstract

Developmental competence allows tissues to respond to inductive cues before committing to specialized forms, but how this potential is encoded at clustered gene-family loci is poorly understood. We use vertebrate skin to address this problem. Epidermis responds to regional dermal signals before committing to feather, scale, or differentiated programs, and α-keratin loci provide a stringent genomic test: separated type-I/type-II clusters show coordinated transcriptional pairing, yet individual keratin genes are selectively deployed across appendage, differentiation, and disease states. Using chicken developmental genomics with comparative mouse and human epidermal datasets, we show that α-keratin clusters are organized before commitment as scaffolded chromatin domains. Within these domains, regulatory elements remain broadly accessible but acquire state-specific activity during commitment and differentiation. Inter-cluster contacts and chromatin-factor perturbation link this architecture to keratin output and morphology. These findings reveal a locus-level chromatin basis for developmental competence, enabling domain-level coordination with gene-level selectivity during epidermal diversification.

## INTRODUCTION

Developmental competence is the capacity of a tissue to interpret inductive signals before its fate becomes restricted. This capacity is transient: a signal can redirect development only while the responding tissue remains competent^1,2^. Skin has provided one of the clearest experimental demonstrations of this principle. In epithelial-mesenchymal recombination studies, regional dermis could redirect epidermal appendage fate only within a finite developmental window, while the epidermis retained the capacity to generate alternative integumentary structures^2–6^. Dermal signals redirected keratin synthesis, epithelial gene networks, and appendage morphology^7,8^, showing that competence includes readiness to execute alternative structural programs. These experiments defined competence at the level of tissue interactions. How such developmental potential is maintained at the genome level remains largely unknown.

This question is especially relevant for clustered gene families. Related genes must often be coordinated, but individual family members must also respond selectively to developmental stage, cellular state, anatomical position, or physiological demand^9,10^. Three-dimensional genome architecture can reconcile these requirements by partitioning chromosomes into regulatory neighborhoods and establishing preferred enhancer-promoter interactions^11,12^. Different clustered loci have evolved distinct architectural solutions. Hox clusters coordinate positional and temporal programs through progressive chromatin-domain transitions and directional insulation^13,14^. Protocadherin clusters use CTCF/cohesin-dependent looping to select among alternative promoters^15^. Olfactory receptor loci assemble interchromosomal enhancer hubs that support singular receptor choice^16^, whereas globin loci use locus-control-region contacts to switch gene expression during development^17^. These systems explain how clustered genes encode positional identity, stochastic choice, or temporal transitions. They do not explain how a structural gene family preserves coordinated output while remaining available for multiple region- and state-specific programs.

The α-keratin family poses this challenge in an unusually stringent form. Type-I and type-II α-keratin genes reside in separate clusters on different chromosomes^18,19^, yet their transcription is coordinated in stratified epidermis^20^. At the same time, the clusters do not behave as uniform expression blocks. Individual keratin genes are selectively deployed across epidermal layers^20,21^, anatomical regions, appendages^22^, differentiation states, injury responses, cancers, and skin diseases^23^. The relevant regulatory problem is therefore not simply how keratin genes are activated. It is how two physically separated gene families maintain coordinated behavior while permitting different keratin combinations to be chosen in different developmental contexts. This requirement distinguishes α-keratin loci from classical cluster-regulation paradigms.

Previous studies have defined individual keratin promoters, differentiation-associated keratin expression, and appendage-specific programs^24^. Regional appendage identity depends on interacting regulatory modules^25^, nuclear architecture contributes directly to epidermal structural-gene regulation. In birds, β-keratin clusters use enhancer-driven chromatin looping to generate region-specific outputs^26^, while SATB2, a nuclear matrix-associated chromatin organizer, shifts α- and β-keratin expression at the cluster level^27^. In mammals, p63 induces SATB1 to reorganize the epidermal differentiation complex and activate terminal differentiation genes^28,29^. Tet2/3-dependent DNA demethylation also shapes chromatin accessibility and three-dimensional organization at lineage-specific epidermal loci^30^. These findings establish that chromatin architecture participates in epidermal differentiation. However, they leave unresolved a fundamental issue of developmental timing: is the regulatory organization of α-keratin loci constructed as keratin programs become active^10,31^, or is it established earlier, while the epidermis still retains alternative developmental options^32,33^?

We address this question by using the shifted developmental timing of chicken skin. Dorsal epidermis enters the feather-forming program earlier, whereas shank epidermis retains competence longer before committing to scale formation^3,5,6^. This temporal offset allows α-keratin regulation to be examined across competence and commitment rather than only between terminal states. We integrate this *in vivo* developmental series with mouse and human epidermal datasets to test whether the resulting regulatory logic is conserved across vertebrate differentiation. Our study reveals that competence is reflected at the level of gene-cluster architecture: α-keratin loci are prepared before commitment so that later programs can select individual genes without rebuilding family-level organization. This identifies a chromatin mechanism by which developing tissues preserve alternative structural outputs while remaining poised for diversification.

## RESULTS

### Conserved α-keratin clusters undergo a competence-to-commitment expression switch

Comparative analysis of α-keratin regulation first required a reliable genomic framework. We used current avian keratin annotations^34^ and reciprocal-hit orthology together with synteny construction to distinguish one-to-one orthologs from lineage-specific one-to-many duplication relationships across chicken, mouse, and human (Figure 1A; Table S1). This registry showed that type-I and type-II α-keratin genes remain clustered across amniotes, despite species-specific changes in gene content, local order, and duplication history. Figure 1A and Figure S1A therefore define the coordinate system for downstream analyses of keratin expression, chromatin architecture, candidate regulatory elements, and inter-cluster contacts.

**Figure 1.**
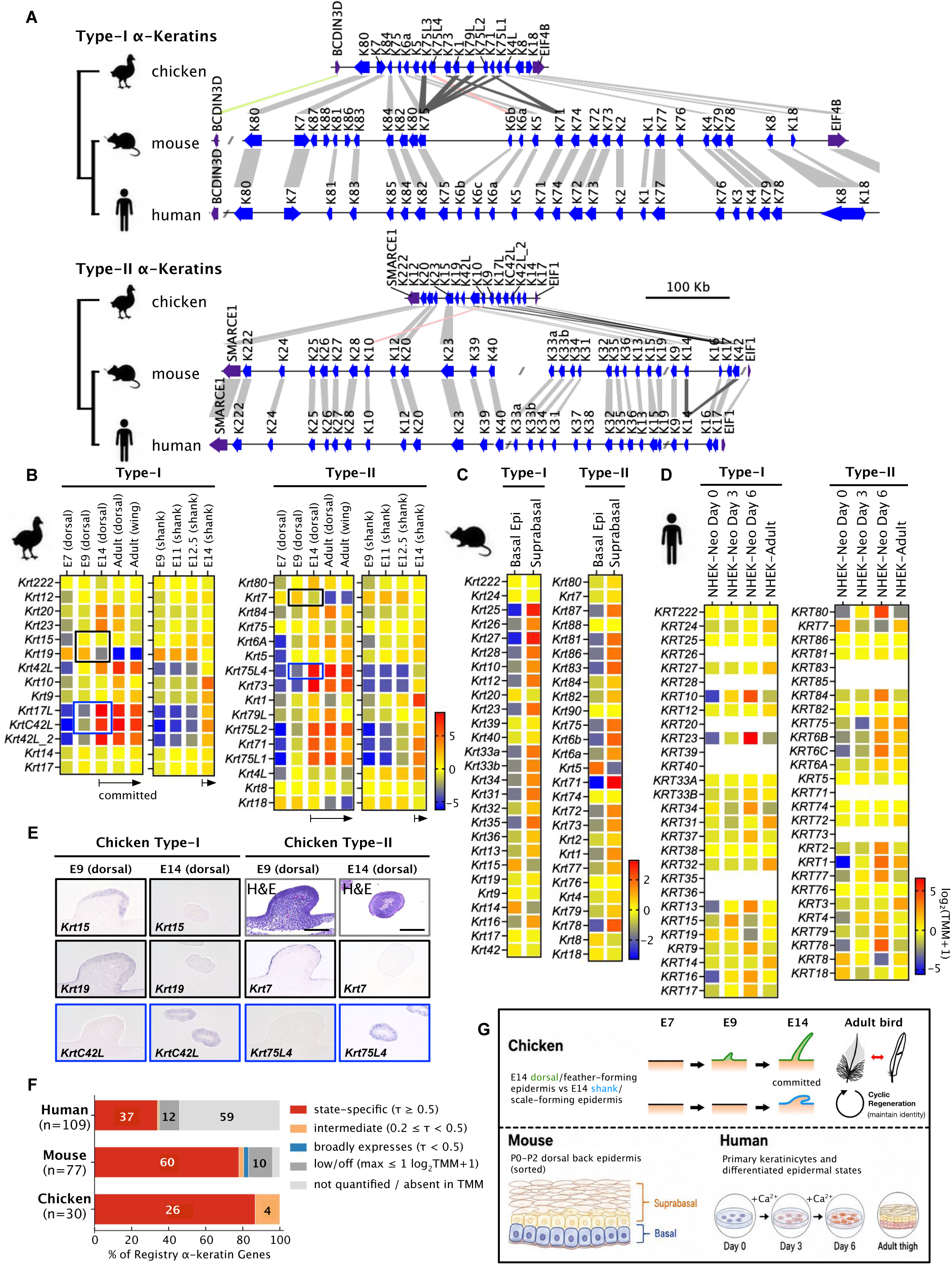
Conserved α-keratin gene clusters support flexible developmental and regional expression programs. See also Figure S1 and Tables S1 and S2. **(A)** Comparative synteny of type-I and type-II α-keratin gene clusters in chicken, mouse, and human. Blue arrows indicate α-keratin genes, and purple arrows indicate neighboring non-keratin genes used to orient the genomic neighborhoods. Connecting lines indicate orthologous or lineage-remodeled relationships: gray, one-to-one orthologs; dark gray, duplication relationships; pink, translocation; green, inversion. Scale bar, 100 kb. **(B–D)** Expression profiles of type-I and type-II α-keratin genes in chicken, mouse, and human epidermal states. Heatmap values represent gene-centered state-mean log₂(TMM + 1) expression calculated separately within each species. Values indicate expression above or below each gene’s own across-state mean and are not absolute expression levels across genes or species. **(B)** Chicken dorsal, feather, and shank/scale epidermal states across embryonic and adult stages. **(C)** Sorted basal and suprabasal keratinocytes from P0–P2 mouse dorsal back skin. **(D)** Human neonatal foreskin keratinocytes at undifferentiated day 0 and calcium-induced differentiation days 3 and 6, together with adult keratinocyte/epidermal states. **(E)** *In situ* hybridization of representative chicken type-I keratins (*Krt15*, *Krt19*, and *KrtC42L*) and type-II keratins (*Krt7* and *Krt75L4*) in E9 and E14 dorsal epidermis. H&E staining is shown for tissue morphology. Black borders indicate genes with higher RNA-seq expression at E9 than E14, and blue borders indicate genes with higher expression at E14 than E9 in the chicken dorsal epidermal series. Scale bar, 100 μm. **(F)** Yanai τ-based classification of α-keratin expression breadth across each species-specific epidermal series. Bars show the percentage of curated registry entries classified as state-specific, intermediate, broadly expressed, low/off, or not quantified/absent in the TMM matrix. Numbers inside bars indicate gene counts. Mouse classifications reflect basal–suprabasal bias because only two mouse epidermal states were analyzed. **(G)** Skin tissues used in this study to demonstrate different developmental stages, anatomical regions, and epidermal differentiation states from three species.

We next asked how this conserved clustered organization is used during epidermal competence, commitment, and differentiation. Chicken provided the developmental series. In the dorsal trajectory, E7 dorsal epidermis represents an early competence stage, E9 dorsal epidermis marks progression into the feather-forming trajectory, E14 dorsal feather epidermis represents committed feather-forming epidermis, and adult feather epidermis provides a mature appendage state. In the shank trajectory, E9, E11, and E12.5 shank epidermis capture the later scale-forming trajectory before full commitment, whereas E14 shank epidermis represents committed scale-forming epidermis. RNA-seq revealed distinct pre-commitment, committed, and mature appendage-associated expression profiles across these feather-forming and scale-forming trajectories (Figure 1B).

To ask whether this flexibility is a broader vertebrate feature, we analyzed mouse^35,36^ and human epidermal datasets^32,37^ (Figure 1C and 1D; Table S2). Mouse basal and suprabasal epidermal cells separated canonical basal keratins, including *Krt5*, *Krt14*, and *Krt15*, from suprabasal keratins such as *Krt1* and *Krt10*. Human keratinocytes showed distinct keratin subsets during calcium-induced differentiation and in adult epidermal states. These mammalian datasets do not provide the same competence-to-commitment timing as chicken embryonic skin, but they show that α-keratin clusters are not deployed as uniform transcriptional blocks across epidermal states.

To compare preferential keratin use across states, we summarized replicate expression within each state as state-mean log₂(TMM + 1) expression^38^ and plotted each gene relative to its own average expression within the same species. This approach highlights state-biased keratin deployment without implying direct comparison of absolute expression across genes or species. In chicken, the key pattern was not simply developmental age. Keratin expression changed as dorsal epidermis progressed from competence toward feather commitment and as shank epidermis progressed toward scale commitment. For example, E14 dorsal feather epidermis showed increased expression of feather-associated type-I keratins, including *Krt17L*, *KrtC42L*, and *Krt42L_2*, while *Krt19* was reduced relative to earlier dorsal stages (Figure 1B). The shank trajectory followed a distinct scale-associated program, separating committed E14 shank epidermis from earlier shank stages and from the dorsal feather program.

We then connected the chicken RNA-seq profiles to tissue-level expression. *In situ* hybridization for representative type-I and type-II keratins showed stage- and region-associated differences between E9 and E14 dorsal epidermis (Figure 1E). *Krt15*, *Krt19*, *KrtC42L*, *Krt7*, and *Krt75L4* therefore support the interpretation that bulk expression differences reflect organized epidermal programs rather than uniform changes across the tissue. Finally, Yanai τ specificity scores showed that keratin genes span state-biased, intermediate, broadly expressed, low/off, and unquantified categories (Figure 1F). Because the mouse comparison contains only basal and suprabasal states, mouse τ values were interpreted as basal-suprabasal bias rather than multi-state specificity.

Figure 1 establishes the developmental problem. Type-I and type-II α-keratin clusters are conserved, but keratin output is not uniform. In chicken, keratin expression separates competence, progression, committed feather, committed scale, and mature feather states. In mouse and human, keratin genes distinguish epidermal differentiation states. Thus, the same clustered gene-family organization supports selective keratin deployment during epidermal differentiation (Figure 1G)

### Keratin-domain architecture is present during epidermal competence before regional appendage commitment

We next asked whether α-keratin chromatin architecture appears mainly when keratin genes become highly active, or whether domain organization is already present before maximal regional keratin deployment. Chicken provided the clearest developmental test because Micro-C was available across dorsal epidermal competence, feather-forming progression, committed feather epidermis, shank/scale epidermis, and adult feather epidermal states. At both type-I and type-II α-keratin loci, local topologically associating domain (TAD) organization was already evident in E7 dorsal epidermis, before maximal feather-associated keratin expression (Figure 2A and Video S1). By E9, as dorsal epidermis progressed into the feather-forming trajectory, internal subdomain organization became more apparent. These domains persisted in E14 dorsal feather epidermis, E14 shank/scale epidermis, and adult feather epidermis. Thus, keratin-domain architecture is not simply a late consequence of maximal keratin transcription. The loci are already organized during competence.

**Figure 2.**
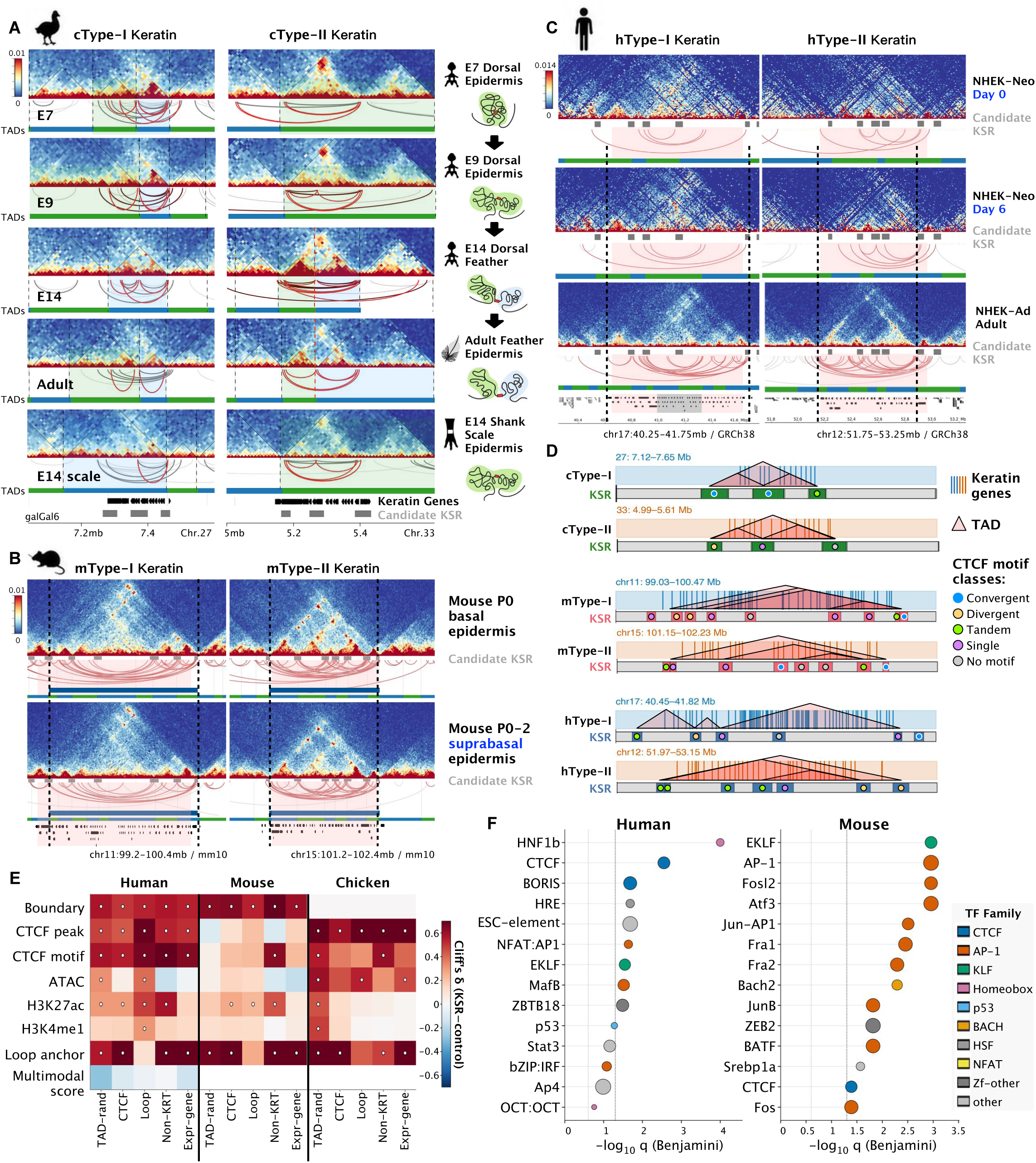
Candidate Keratin Scaffold Regions mark conserved architectural domains at α-keratin loci. See also Figure S2. **(A)** Micro-C contact maps of chicken type-I and type-II á-keratin loci across developmental and regional epidermal states, including early dorsal epidermis, feather-forming epidermis, shank/scale-forming epidermis, and adult feather epidermis. Arcs indicate called chromatin contacts; RNA-seq tracks show keratin expression across the same loci. **(B)** Hi-C contact maps of mouse type-I and type-II keratin loci in sorted P0–P2 basal and suprabasal epidermal cells. Arcs indicate called chromatin loops, and RNA-seq tracks show matched keratin expression. **(C)** Contact maps of human type-I and type-II keratin loci in neonatal keratinocytes during calcium-induced differentiation and in adult keratinocytes. Arcs indicate called loops, and RNA-seq tracks show keratin expression across differentiation states. **(D)** Cross-species schematic of candidate Keratin Scaffold Regions (candidate KSRs) at human, mouse, and chicken type-I and type-II α-keratin loci. Candidate KSRs mark boundary-associated regions that flank or subdivide keratin chromatin domains. **(E)** Matched-control analysis comparing active candidate KSRs with within-TAD random intervals, CTCF-anchor controls, loop-anchor controls, non-keratin cluster boundaries, and expressed-gene boundaries. Color indicates Cliff’s ä, and dots mark comparisons passing both BH-FDR and effect-size thresholds. Human KSRs show the strongest separation from CTCF- and loop-anchor controls; mouse and chicken show more feature- and context-dependent separation. **(F)** Motif enrichment at accessible regions within active candidate KSRs. HOMER known-motif enrichment was performed using ATAC peaks overlapping candidate KSRs as foreground and all genome-wide ATAC peaks as background. Dot position indicates −log10 Benjamini q value, dot size indicates the fraction of foreground peaks containing the motif, and color indicates TF family. Chicken motif discovery was underpowered because of the smaller foreground set.

We then asked whether this organization is restricted to chicken. In mouse basal and suprabasal epidermal cells, Hi-C maps showed that type-I and type-II keratin loci are organized within local chromatin domains (Figure 2B). Human neonatal keratinocytes undergoing calcium-induced differentiation, together with adult keratinocyte datasets, showed comparable domain organization and loop structure at the corresponding α-keratin loci (Figure 2C). These mammalian datasets do not provide the same developmental timing as chicken embryonic skin, but they show that keratin-domain architecture is retained across vertebrate epidermal states.

To define landmarks within these domains, we avoided simply naming visual boundaries, generic CTCF sites, loop anchors, or active chromatin near highly expressed genes. We defined candidate Keratin Scaffold Regions (KSRs) using chromatin topology as the primary criterion. Candidate intervals were called from contact-derived boundary structure within keratin-domain discovery windows and then annotated for CTCF ChIP-seq or motif support, ATAC accessibility, active histone marks, and loop/contact context (Figure S2A). In this design, contact maps define candidate KSRs; chromatin and motif features test whether these contact-derived intervals carry expected architectural or regulatory features.

This approach identified 36 active candidate KSRs across the three species: 13 in human, 17 in mouse, and 6 in chicken (Figure 2D; Table S3). Candidate KSRs were present at both type-I and type-II loci in each species. Most were supported by multiple evidence layers, including contact-boundary structure, CTCF annotation, and accessible or active chromatin. They were not restricted to promoters, but occurred at domain-flanking regions and internal sub-boundaries, consistent with landmarks that demarcate and subdivide α-keratin domains (Figure S2B).

We next tested whether candidate KSRs could be explained by generic architectural or active-locus features. Candidate KSRs were compared with matched within-TAD random intervals, CTCF-anchor controls, loop-anchor controls, non-keratin cluster boundaries, and expressed-gene boundaries (Figure 2E). Human KSRs showed the strongest separation, retaining five of nine tested features against CTCF-anchor controls and six of nine against loop-anchor controls after multiple-testing and effect-size filtering (Figure S2C). Mouse and chicken showed more feature-dependent separation, likely reflecting mixed-context mouse CTCF data and the smaller number of chicken KSRs. These comparisons support candidate KSRs as keratin-locus architectural candidates, but do not justify treating KSRs as a new genome-wide class of architectural elements. CTCF motif orientation provided an additional but limited test. Convergent orientation was not the dominant pooled motif class across candidate KSRs (Figure S2D and S2E), indicating that KSRs are not simply generic convergent CTCF sites.

Accessible regions within candidate KSRs carried sequence features consistent with architectural and epidermal regulatory inputs. Using ATAC peaks overlapping active candidate KSRs as foreground and genome-wide ATAC peaks as background, motif enrichment recovered CTCF and KLF-family motifs in human and mouse, with AP-1/bZIP-family enrichment most prominent in mouse (Figure 2F). Pairing enriched motifs with transcription-factor expression helped distinguish plausible motif-TF relationships from unresolved motif assignments (Figure S2F). Chicken motif discovery was underpowered because only 49 KSR-overlapping accessible peaks were available, so no chicken-specific motif claim was made.

The unexpected result from the chicken Micro-C series was that α-keratin domains were already visible at E7 dorsal epidermis, before maximal feather- or scale-associated keratin deployment. Later stages did not create the architecture *de novo*; they refined an existing domain structure as epidermis progressed toward feather and scale commitment. Candidate KSRs mark contact-derived borders and internal sub-boundaries within this competence-associated scaffold. Figure 2 therefore places α-keratin-domain architecture before maximal regional keratin activation and identifies candidate landmarks that organize the loci.

### Broadly accessible regulatory elements acquire state-specific activity within keratin-domain scaffolds during keratin deployment

Having shown that α-keratin loci are already organized during competence, we next asked how individual keratin genes are selected within this scaffold. If regional keratin programs are driven mainly by activation-coupled regulatory remodeling, accessible regulatory elements should appear primarily when feather-, scale-, basal-, suprabasal-, or differentiated keratin programs become active. If keratin domains are already competent, many regulatory elements should be accessible earlier and later acquire state-specific activity or contact context.

We analyzed chromatin accessibility, histone modifications, CTCF occupancy, and contact context across type-I and type-II α-keratin loci, combining chicken developmental skin profiles with published mouse and human epidermal datasets where available (Table S2). Across all three species, many promoter-proximal and distal accessible elements were present within keratin domains before or across major changes in keratin expression (Figure 3A–D; Figure S3A–I). In chicken, accessible elements were detected in early dorsal epidermis and persisted into later feather- and scale-forming states. Published mouse basal/suprabasal epidermal datasets and human keratinocyte differentiation datasets showed similar retention of accessible elements across epidermal-state transitions. Thus, keratin deployment is not explained by late opening of an otherwise inaccessible locus.

**Figure 3.**
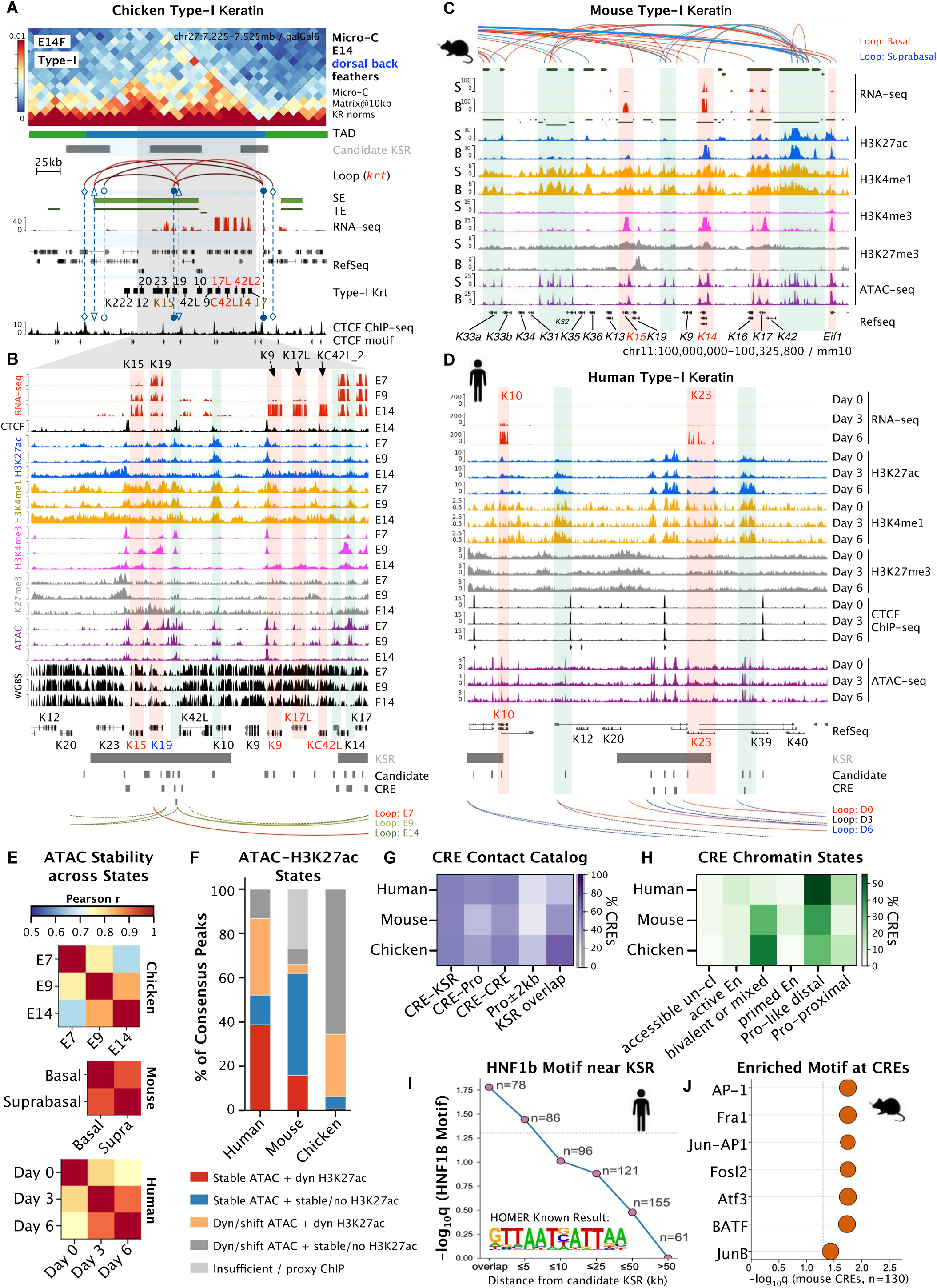
Accessible regulatory landscapes within KSR-defined domains accompany gene-specific keratin output. See also Figure S3. **(A and B)** Developmental multi-omic views of chicken α-keratin loci across dorsal epidermis, feather-forming epidermis, shank/scale-forming epidermis, and adult feather epidermis. Tracks show RNA-seq, histone modifications, ATAC-seq, DNA methylation, CTCF ChIP-seq, and Micro-C contacts. Candidate KSRs and KSR-bounded intervals are indicated. **(C and D)** Comparative mammalian multi-omic views of keratin loci in mouse P0 basal and suprabasal epidermal cells (C) and human keratinocytes during calcium-induced differentiation (D). These profiles show established keratin-domain regulatory landscapes in mammalian epidermal states. **(E)** ATAC signal similarity across matched keratinocyte or epidermal states. Heatmaps show Pearson correlations of state-mean ATAC signal over consensus keratin-domain peaks in chicken dorsal epidermis-to-feather trajectory, mouse basal versus suprabasal epidermis, and human NHEK differentiation. **(F)** ATAC–H3K27ac classes at candidate CREs. Stacked bars show the distribution of candidate CREs across five accessibility–activity classes in human, mouse, and chicken. Classes summarize whether ATAC accessibility is stable or shifted across stages and whether H3K27ac is stable/absent or dynamic. “Insufficient/proxy ChIP” denotes missing stage-matched H3K27ac data or proxy tracks. **(G)** Contact and overlap support for candidate CREs. Grouped bars show the fraction of CREs supported by CRE–KSR contact, CRE–CRE contact, CRE–promoter contact, KSR overlap, or promoter proximity. CRE–KSR contact support is the most consistent contact-associated feature across species. **(H**) Chromatin-state classification of candidate CREs. Bars show the fraction of CREs assigned to promoter-like distal, promoter-proximal, active-enhancer-like, primed-enhancer-like, bivalent/mixed, repressed/polycomb-associated, or accessible-unclassified states. Human CREs are enriched for promoter-like distal states, whereas mouse and chicken show larger bivalent/mixed fractions. **(I)** Human HNF1b motif enrichment as a function of CRE distance from active candidate KSRs. Points show −log10 Benjamini q values across KSR-distance bands, with foreground size indicated for each band. **(J)** Mouse AP-1/bZIP motif enrichment at candidate CREs. Dot plots show AP-1 family motif enrichment in mouse CREs; dot size indicates the fraction of target sequences containing the motif. Corresponding AP-1 family TFs are expressed across mouse keratinocyte states. Motif results nominate candidate TF families but do not establish TF occupancy or regulatory function.

To quantify this behavior, we built consensus ATAC peak sets within keratin domains and measured accessibility similarity across matched epidermal stages or cell states (Figure 3E; Figure S3J–L). Normal human epidermal keratinocyte (NHEK) D0, D3, and D6 showed highly similar ATAC profiles across 286 consensus peaks, with mean pairwise Pearson *r* = 0.819. Mouse P0 basal and suprabasal epidermal cells showed comparable stability across 226 peaks, with r = 0.799. In chicken, the matched dorsal trajectory— E7 dorsal epidermis, E9 dorsal epidermis, and E14 dorsal feather filament—showed stronger internal ATAC similarity than the mixed dorsal-plus-leg comparison, with r = 0.760 versus 0.716. This dorsal trajectory also identified 40 of 145 peaks with early or persistent accessibility. These results indicate that keratin-domain accessibility is broadly retained across related epidermal trajectories rather than established only at committed expression states.

Stable accessibility did not mean stable regulatory activity. Integrating ATAC with H3K27ac showed that subsets of accessible elements retained accessibility while changing enhancer-associated activity marks. In human and mouse, 56 of 286 human peaks and 36 of 226 mouse peaks showed stable ATAC with dynamic H3K27ac (Figure 3F). In these mammalian datasets, accessibility stability was similar to matched non-keratin backgrounds, suggesting that pre-accessibility is a broader feature of epidermal chromatin rather than a keratin-exclusive property. In chicken, keratin-domain ATAC was more stable than matched non-keratin background, with mean signal coefficient of variation (CV) of 0.50 versus 0.97, although this comparison remains limited by sample structure and foreground size.

To organize this regulatory landscape, we constructed an ATAC-seeded candidate cis-regulatory element (CRE) catalog within α-keratin domains. This catalog contained 406 candidate CREs: 216 in human, 130 in mouse, and 60 in chicken (Figure 3G; Table S4). Candidate CREs included promoter-proximal elements, distal non-promoter elements, and a KSR-overlapping class. Across species, CRE– KSR contact support was the most reproducible contact-associated feature, accounting for 58–62% of cataloged CREs. This suggests that many candidate regulatory elements lie within or near the architectural scaffold rather than acting only from isolated promoter-proximal positions.

We classified candidate CRE states using H3K27ac, H3K4me1, H3K4me3, H3K27me3, and proximity to canonical keratin transcription start sites (Figure 3H). Human candidate CREs were dominated by promoter-like distal elements, whereas mouse and chicken contained larger bivalent or mixed fractions. Because the available datasets differ in stage and tissue context—differentiating and adult human keratinocytes, mouse P0 epidermis, and embryonic chicken skin—we interpret these differences descriptively rather than as fixed species-specific states. Motif enrichment was used only as a prioritization layer. It highlighted a human *HNF1b* proximity-decay signal near active KSRs, ZEB2 enrichment at mouse CRE–KSR-loop-associated elements, broad AP-1-family enrichment in mouse CREs, and exploratory Hox/RUNX1 enrichment at chicken bivalent CREs (Figure 3I and 3J; Figure S3O and S3Q; Table S5). These motifs nominate candidate TF families but do not establish TF occupancy or causal regulation.

Figure 3 shows that accessibility alone cannot explain keratin-gene choice. Many α-keratin-domain elements are already open across related epidermal states, whereas H3K27ac, promoter context, KSR proximity, contact support, and motif-associated features vary more selectively. The regulatory transition is therefore not wholesale opening of a closed locus. It is differential use of a pre-accessible landscape within an already organized keratin domain.

### Inter-cluster coordination and chromatin-factor perturbation link keratin architecture to epidermal morphology

Local domain architecture explains how each α-keratin cluster is organized, but it does not fully solve the paired-gene-family problem. Type-I and type-II keratin genes reside on separate chromosomes, yet stratified epidermis deploys coordinated type-I/type-II transcriptional programs. We therefore asked whether separated type-I and type-II domains show higher-order population-level proximity during keratin deployment, and whether perturbing chromatin-associated factors alters keratin output and epidermal morphology.

In chicken E14 feather filaments, FitHiC2 prioritized contacts^39^ between the type-I cluster on chromosome 27 and the type-II cluster on chromosome 33 (Figure 4A; mouse comparison in Figure S4A). These contacts were concentrated near candidate contact-prioritized regions positioned at TAD boundaries, candidate KSRs, and putative super-enhancer intervals (Figure 4B and Figure S4B). Because focal interchromosomal pixels are sensitive to sequencing depth, sparsity, and normalization, we treated these regions as candidate landmarks and tested the higher-order signal at the domain level using matched genomic backgrounds.

**Figure 4.**
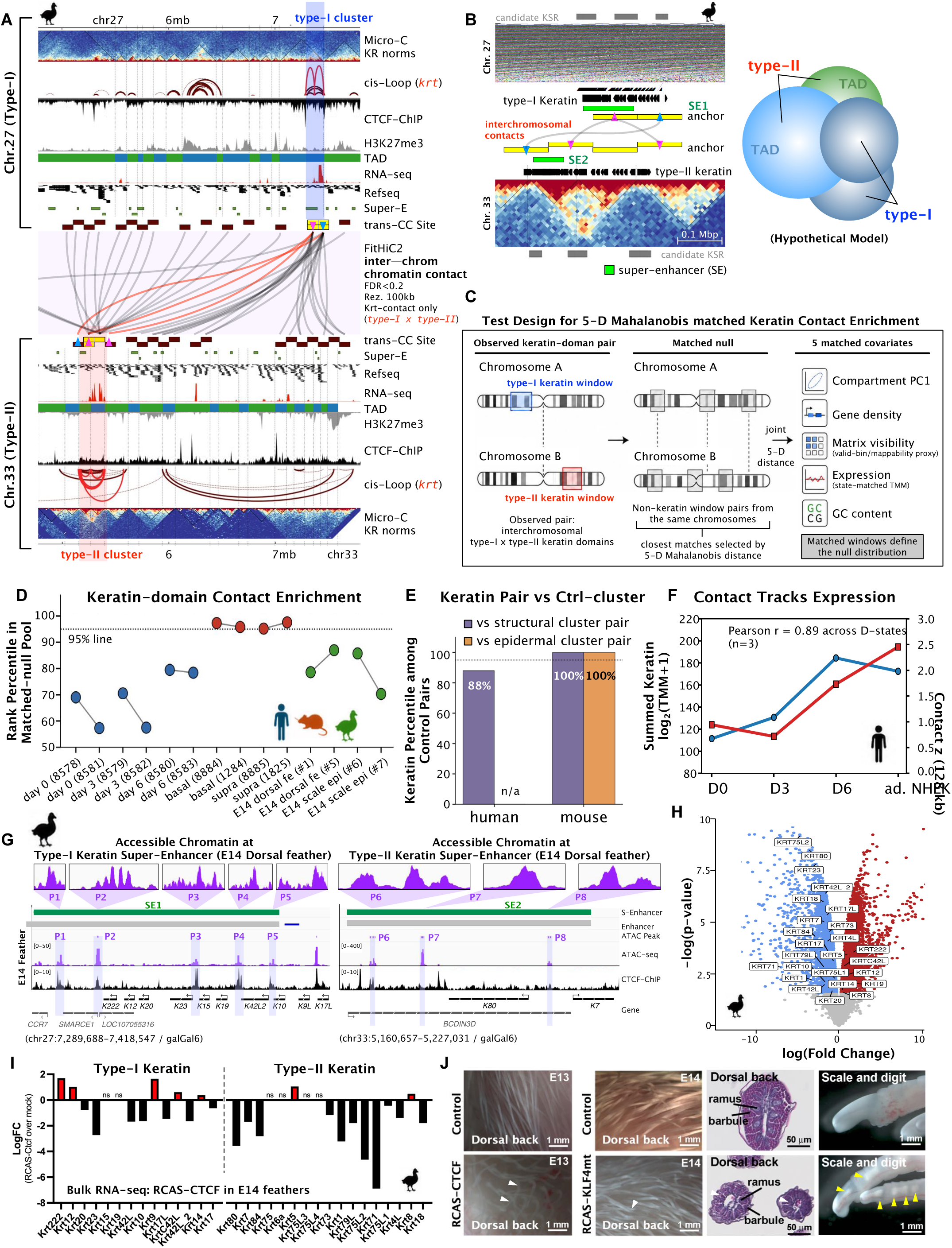
Higher-order keratin-cluster organization and chromatin perturbation link architecture to epidermal function. See also Figure S4. **(A)** FitHiC2 results showing interchromosomal contacts between the type-I keratin cluster on chicken chromosome 27 and the type-II keratin cluster on chromosome 33. Red lines indicate significant trans contacts. RNA-seq tracks show normalized read coverage in bins per million mapped reads. ChIP-seq tracks show fold enrichment over input. The chicken genome assembly used is GRCg6a. **(B)** Summary of inter-cluster contacts between type-I and type-II keratin loci, highlighting contact anchors associated with TAD boundaries, KSRs, and super-enhancer regions. **(C)** Five-covariate matched-null test design. The observed type-I × type-II α-keratin window pair was compared with non-keratin window pairs drawn from the same chromosomes and matched by Mahalanobis nearest-neighbor search across five covariates: A/B-compartment PC1, gene density, state-matched TMM expression, GC fraction, and a mappability proxy based on matrix visibility. For each sample, 200 nearest non-keratin windows were selected per chromosome, yielding 40,000 matched non-keratin window pairs at 128-kb contact resolution. Empirical *p*-values were calculated as the fraction of matched-null pairs with balanced contact greater than or equal to the observed keratin-domain contact; rank percentiles were assigned 95% confidence intervals by 1,000 bootstrap iterations. **(D)** Replicate-resolved keratin-domain contact enrichment. Per-sample rank percentile of the observed type-I × type-II α-keratin contact within the five-covariate Mahalanobis matched-null distribution for seven biological-state replicate pairs: human D0, D3, and D6; mouse P0 basal and P0 suprabasal; chicken E14 feather and E14 scale. Dots indicate rank percentile; whiskers indicate 95% bootstrap confidence intervals from 1,000 iterations; grey lines connect biological replicates within each state. The dotted line marks the 95th percentile. **(E)** Keratin-domain contact enrichment relative to control gene-cluster pairs. Percentile rank of the observed type-I/type-II α-keratin contact relative to matched non-keratin gene-cluster controls. A matched cross-chromosome active-epidermal control pool could not be constructed for human because the major epidermal differentiation complex resides on chr1 and did not provide eligible cross-chromosome matched pairs in this analysis. **(F)** Contact enrichment tracks keratin expression. Dual-axis plots compare summed α-keratin expression, shown as log₂(TMM + 1), with type-I/type-II trans-contact z score across matched epidermal states. Human neonatal keratinocytes are shown across D0, D3, and D6, with adult NHEK included as an additional state. Human D0–D6 states show a positive association between keratin expression and trans-contact z score (Pearson *r* = 0.89, n = 3). Adult keratinocytes are shown descriptively. **(G)** ATAC-seq and CTCF ChIP-seq profiles at putative super-enhancer regions associated with the type-I and type-II keratin loci. ATAC-seq tracks show normalized pileup values calculated by Genrich. SE, super-enhancer; TE, typical enhancer. **(H-I)** Differential expression analysis of E14 feather epidermis following RCAS-mediated CTCF overexpression, highlighting altered expression of α-keratin genes. **(J)** Epidermal phenotypes following CTCF overexpression and KLF4 mutant overexpression in E14 chicken embryos, including abnormal feather filament morphology (white arrows) and reduced keratinization in digit skin. Notably, KLF4 mutant overexpression rendered the digit skin transparent (yellow arrows).

Keratin loci are gene-rich, transcriptionally active, and embedded in active chromatin (Figure S4C), any of which could elevate interchromosomal contact frequency. We therefore compared each observed type-I/type-II keratin-domain pair with non-keratin window pairs drawn from the same chromosome pairs and jointly matched for compartment PC1, gene density, state-matched TMM expression, GC content, and matrix-visibility-based mappability using five-dimensional Mahalanobis nearest-neighbor matching^40^ (Figure 4C). This matched-null analysis asked whether type-I/type-II keratin-domain proximity exceeds expectation for active genomic regions with similar chromatin context and technical visibility.

Mouse epidermis provided the strongest evidence for higher-order type-I/type-II keratin-domain proximity. In the replicate-resolved analysis, all four mouse basal and suprabasal epidermal samples exceeded the 95th-percentile matched-null threshold, with rank percentiles of 95.2%-97.6% and empirical *p*-values of 0.024-0.048 (Figure 4D). Human keratinocytes did not cross the significance threshold but showed a differentiation-associated directional increase, with D6 replicates ranking higher and more concordantly than D0 or D3 replicates. Chicken E14 feather and scale samples were also directionally elevated but not significant. The full 21-sample analysis preserved this hierarchy, with mouse epidermal samples ranking highest, human samples intermediate, chicken epidermal or appendage samples generally above chicken brain controls, and the lowest chicken rank observed in E14 brain (Figure S4E). Thus, domain-level type-I/type-II proximity is statistically supported in mouse epidermis, directional during human keratinocyte differentiation, and suggestive in chicken appendage contexts.

We next asked whether this proximity was merely a general property of active gene clusters. In mouse, the type-I/type-II α-keratin pair exceeded both matched structural gene-cluster controls and active epidermal gene-cluster controls, ranking at the 100th percentile in each comparison (Figure 4E). In human, the keratin pair ranked at the 88th percentile relative to structural gene-cluster controls, although an active epidermal cross-chromosome control pool could not be constructed. The mouse signal was robust to covariate removal: in P0 basal epidermis, the keratin-pair rank remained between 96.77% and 97.72% after removing any single matching covariate (Figure S4F). In human neonatal keratinocytes, type-I/type-II contact z scores increased with summed α-keratin log₂(TMM + 1) expression across D0, D3, and D6, with Pearson *r* = 0.89 (Figure 4F). Mouse basal and suprabasal states showed the same direction. Because pixel-level focal contacts were less reproducible than the domain-level signal, candidate contact-prioritized intervals are reported as a follow-up resource in Table S6 rather than used as the primary evidence for higher-order proximity.

We returned to chicken E14 feather loci to connect candidate higher-order interaction zones with the local regulatory landscape. Candidate interaction zones were concentrated near KSRs and putative super-enhancer intervals (Figure 4A and 4B). At both type-I and type-II α-keratin loci, these candidate super-enhancer regions showed chromatin accessibility and CTCF occupancy (Figure 4G). Motif analysis connected these regions with sequence features identified earlier at candidate KSRs and CREs, including CTCF and KLF-family motifs (Figures 2F, 3I–J, and S4H). KLF4 has been linked to epithelial differentiation and specialized keratin expression in WNT10A-dependent ectodermal programs^41^. Thus, chicken feather epidermis links candidate higher-order interaction zones to the architectural and regulatory features that organize local keratin domains.

Because proximity alone cannot establish biological relevance, we next asked whether perturbing chromatin-associated regulatory activity affects keratin transcription and epidermal form. RCAS-mediated CTCF overexpression in E14 feather epidermis altered α-keratin gene expression across both type-I and type-II clusters (Figure 4H and 4I). Perturbation of CTCF- or KLF4-associated regulatory activity also produced epidermal defects, including abnormal feather filament morphology and disrupted scale organization (Figure 4J). These experiments do not assign individual KSRs, candidate super-enhancers, or contact-prioritized regions to specific target genes, and they do not prove that type-I/type-II proximity is causal. They do show that chromatin-associated regulatory factors can influence keratin transcriptional output and epidermal morphology.

Figure 4 extends the local scaffold to the paired-cluster problem. Separated type-I and type-II domains show population-level proximity, strongest in mouse epidermis and directional in human and chicken contexts. Chromatin-factor perturbation alters keratin expression and epidermal morphology. Thus, paired keratin deployment is associated with higher-order domain coordination and is sensitive to chromatin-associated regulatory state.

## DISCUSSION

Our central finding is that the genome prepares epidermal structural programs before the tissue commits to them. α-Keratin domains are already organized during developmental competence, before maximal feather-, scale-, or differentiation-associated keratin expression. Commitment therefore does not build the keratin regulatory landscape *de novo*. It selects from a pre-organized one. This identifies chromatin architecture as a molecular substrate of developmental competence.

This conclusion extends earlier observations that chromatin topology can precede transcription. Stable enhancer-promoter configurations have been detected before gene activation^32^, and preformed topology at the *Shh* locus supports robust transcription during limb development^33^. At α-keratin loci, however, the principle operates across an entire structural gene family. Both type-I and type-II clusters are organized before their state-specific transcriptional programs are fully deployed. The relevant unit of competence is therefore not only an enhancer or promoter, but a gene-cluster domain.

The scaffold does not predetermine one epidermal fate. Many regulatory elements remain accessible across developmental states, whereas H3K27ac, local contact context, and transcription change more selectively. This separation between regulatory availability and regulatory activity is consistent with studies distinguishing poised or accessible elements from active enhancers^42^. It also explains how stable architecture can support changing output. Although this organization is permissive rather than fate-determining, disruption of domain organization can rewire enhancer–gene communication^43^. We propose that the domain preserves regulatory availability, while commitment-associated factors select among accessible elements and keratin genes.

This model resolves the defining problem of the α-keratin family. Type-I and type-II genes occupy separate chromosomes yet support coordinated transcriptional programs, while individual genes are selected across epidermal regions and differentiation states. The loci achieve domain-level coordination with gene-level selectivity. Domain architecture preserves family-level organization, whereas state-specific regulatory activity selects individual keratin outputs. This differs from ordered Hox activation, singular olfactory receptor choice, and globin switching: keratin loci must remain coordinated without losing combinatorial flexibility.

The mammalian epidermal literature supports an architectural role in differentiation. p63-dependent SATB1 activity reorganizes the epidermal differentiation complex^28,29^, while Tet2/3-dependent DNA modification shapes accessibility and three-dimensional organization at lineage-specific loci^30^. Our study adds the missing developmental timing. At α-keratin loci, substantial domain organization is present before commitment, rather than arising only after terminal differentiation begins. Chicken establishes this temporal sequence *in vivo*; mouse and human datasets show that organized domains remain compatible with flexible keratin use across mammalian epidermal states.

Type-I/type-II interchromosomal proximity may provide an additional layer of family coordination. Co-regulated genes can occupy shared transcriptional environments^44^, including preferential associations among erythroid genes^45^. In our datasets, this higher-order association is most pronounced in mouse epidermis and shows the same directional relationship in human and chicken, supporting a shared coordinating principle whose strength varies across species and epidermal states. CTCF- and KLF-associated perturbations further connect chromatin regulation to keratin output and morphology. Together, these findings extend the scaffold model from local domain organization to higher-order coordination, while leaving the specific contacts that mediate this coupling to be resolved.

The broader implication is that developmental competence is not carried only by signaling pathways and transcription factors. It can also reside in the prior organization of the loci that execute tissue form. By architecturally preparing α-keratin clusters before commitment, epidermis preserves alternative structural outputs while remaining able to choose among them. Competence is therefore not simply the ability to receive an inductive signal; it is the genomic capacity to translate that signal into alternative tissue programs.

### Limitations of the study

This study defines an architectural framework rather than a complete causal wiring diagram. Candidate KSRs, CREs, super-enhancer intervals, and contact-prioritized anchors remain unvalidated at the locus level. Hi-C and Micro-C report population-averaged contacts and do not establish single-cell co-localization. CTCF and KLF4 perturbations are global and cannot identify the responsible element or interaction. Targeted editing, Capture-C or 4C, and DNA-FISH will be required to determine how specific features within the scaffold control individual keratin genes.

## Supporting information

Table S5. Candidate CRE motif enrichment and TF-expression integration across three species

## RESOURCE AVAILABILITY

### Lead contact

Further information and requests for resources should be directed to and will be fulfilled by Ya-Chen Liang.

### Materials availability

This study did not generate new unique reagents. Plasmids generated in this study will be made available on request, but we may require a payment and/or a completed materials transfer agreement if there is potential for commercial application.

### Data and code availability

- Newly generated sequencing data, including chicken bulk RNA-seq, ATAC-seq, Micro-C, and WGBS datasets generated for this study, have been deposited in the Gene Expression Omnibus under accession GEO: [pending, reviewer token will be provided] and will be publicly available as of the date of publication.
- Previously published RNA-seq, ATAC-seq, ChIP-seq, Hi-C, and related genomic datasets reanalyzed in this study are listed in Table S2 with accession numbers, source publications, genome builds, and usage annotations.
- Microscopy data reported in this paper will be shared by the lead contact upon request.
- Original analysis code and scripts have been archived through GitHub and Zenodo. The peer-review release of the repository, keratin-cluster-architecture-code v1.0.0-rc1, is available to reviewers through Zenodo at DOI: 10.5281/zenodo.21258190. Reviewer preview link: https://zenodo.org/records/21258190?preview=1&token=eyJhbGciOiJIUzUxMiJ9.eyJpZCI6ImQ4ZjBkMjNmLWFhZjEtNDZjMS05YTY1LWQ2ODVmMjQwYWFlMCIsImRhdGEiOnt9LCJyYW5kb20iOiIxYmIyOTNjNzk1ZTAwYjMyMjg0NTBlNmFiZTYwYTEyZiJ9.o3ITn8tn_Y0HYqQjI5ltJBVrH7NQs2ocCIOD1oRBaO2qvDIUo4MlIFMeu9VG6rsxxerBgqHjnRZkUHZGj6Kb1A
- All data reported in this paper will be shared by the lead contact upon reasonable request.
- Any additional information required to reanalyze the data reported in this paper is available from the lead contact upon reasonable request.

## ACKNOWLEDGMENTS

This work was supported by the National Institute of General Medical Sciences grant R35GM150714 to Y.-C.L. and the National Institute of Arthritis and Musculoskeletal and Skin Diseases grant R01AR047364 to C.-M.C. W.-C.J. is an awardee of the Taiwan-USC Scholarships, a joint partnership between the University of Southern California and the Taiwan Ministry of Education. We thank members of the Chuong and Liang research group for discussions.

## AUTHOR CONTRIBUTIONS

Conceptualization, writing-review & editing, funding acquisition, and supervision: Y.-C.L. and C.-M.C.; methodology, W.-C.J., C.-K.C., and P.W.; Investigation, Y.-C.L., W.-C.J., C.-K.C., and P.W.; writing-original draft, Y.-C.L.

## DECLARATION OF INTERESTS

The authors declare no competing interests.

## DECLARATION OF GENERATIVE AI AND AI-ASSISTED TECHNOLOGIES

During preparation of this work, the authors used ChatGPT, Claude, Grok, and Grammarly to assist with language editing, code organization, and code review. The authors reviewed and verified all resulting content and take full responsibility for the manuscript.

## SUPPLEMENTAL INFORMATION

**Figure S1. Genomic coordinates of type-I and type-II keratin gene clusters in chicken, mouse, and human.**

**Figure S2. Feature annotation and motif support of candidate Keratin Scaffold Regions. Related to Figure 2**.

**(A)** Workflow for annotating contact-derived candidate KSRs. Candidate intervals were centered on local boundary seeds identified from chromatin contact maps and annotated for CTCF ChIP-seq signal, CTCF motif support, ATAC accessibility, H3K27ac/H3K4me1 active chromatin marks, and loop/contact context. Candidate KSRs were classified as multimodal, CTCF-annotated, non-CTCF-annotated, or boundary-only intervals. The KSR definition was based on contact-derived topology; chromatin and motif features were used as supporting annotations.

**(B)** Evidence-layer summary for the 36 active candidate KSRs in the KSRC v1.5G set across human, mouse, and chicken type-I and type-II α-keratin loci. Each symbol represents one candidate KSR. Symbol position indicates relative location within the corresponding keratin cluster, symbol shape indicates position class, and tile stacks indicate support from boundary evidence, CTCF ChIP-seq, CTCF motif, and ATAC accessibility. Candidate KSRs occur at both domain-flanking regions and intra-cluster sub-boundaries.

**(C)** Feature distributions for human active candidate KSRs compared with matched architectural and active-locus controls. Control sets include within-TAD random intervals, CTCF-anchor-matched intervals, loop-anchor-matched intervals, non-keratin cluster boundaries, and expressed-gene boundaries. Features shown include CTCF peak count, CTCF motif count, TAD-boundary support, loop-anchor support, H3K27ac overlap, and ATAC peak count. These comparisons test whether KSRs can be explained as generic CTCF sites, loop anchors, or active-gene boundaries.

**(D)** Schematic of CTCF motif-orientation classes used for KSR annotation. Motif configurations were classified as convergent, divergent, tandem, single motif, mixed, or no motif. Orientation was used as an architectural annotation layer and not as a criterion for defining candidate KSRs.

**(E)** CTCF motif-orientation composition across active candidate KSRs by species and keratin cluster. Bars show the number of KSRs assigned to each motif-orientation class. Convergent orientation was not the dominant pooled class across all KSRs, but provided limited architectural support in specific species or cluster contexts. These data do not establish CTCF-dependent KSR function.

**(F)** Motif logos and TF expression for enriched motifs at accessible candidate KSRs. Rows show enriched HOMER motifs, corresponding position-weight matrix (PWM) logos, and state-mean expression of the queried TF gene across human, mouse, and chicken keratinocyte states. Hatched cells indicate species-specific paralog substitutions used to address annotation gaps, and empty cells indicate unresolved TF assignment. Motif enrichment supports candidate regulatory competence at accessible KSRs, but does not validate TF occupancy or regulatory function.

**Figure S3. Candidate CRE landscapes within KSR-defined α-keratin domains. Related to Figure 3. (A–C)** Chicken developmental multi-omic profiles across type-I and type-II α-keratin loci. Tracks show RNA-seq, histone-modification ChIP-seq, ATAC-seq, DNA methylation, CTCF occupancy, and Micro-C contacts across early dorsal epidermis, feather-forming epidermis, shank/scale-forming epidermis, and adult feather epidermis. Candidate KSRs and KSR-bounded intervals are indicated. These panels show that accessible elements and activity-associated chromatin features are embedded within the KSR-defined architectural framework.

**(D and E)** Hi-C contact maps of mouse type-I and type-II keratin loci in sorted basal and suprabasal epidermal cells from P0–P2 dorsal skin. Loop calls are displayed over the contact maps to show established chromatin organization at the keratin clusters in mammalian epidermal states.

**(D′ and E′)** Matched mouse multi-omic tracks across the same keratin loci, including RNA-seq, histone modifications, ATAC-seq, and CTCF occupancy. These profiles relate keratin expression and chromatin activity to the local contact architecture shown in panels D and E.

**(F and H′)** Human contact maps of α-keratin loci in neonatal keratinocytes during calcium-induced differentiation and in adult keratinocytes. These panels show established chromatin-domain and loop organization at mammalian α-keratin loci.

**(G–I)** Human multi-omic tracks across type-I and type-II keratin loci. Tracks show keratin expression, histone modifications, chromatin accessibility, and CTCF occupancy across differentiation states, providing a mammalian comparison to the developmental chicken and mouse epidermal datasets.

**(J)** Signal-based classification of consensus ATAC peaks within keratin domains. Consensus peaks were classified as stable, quantitatively shifted, or dynamic based on the maximum pairwise ATAC signal change across matched states. This classification summarizes accessibility behavior and does not represent a count-based differential-accessibility test.

**(K)** Chicken dorsal-only ATAC similarity compared with dorsal-versus-leg/scale similarity. ATAC signal was measured over the chicken dorsal consensus peak set to compare similarity within the dorsal epidermis-to-feather trajectory and between dorsal and shank/scale contexts.

**(L)** Comparison of ATAC signal variability between keratin-domain consensus peaks and matched non-keratin background peaks. Human and mouse keratin-domain peaks showed variability similar to matched background regions, consistent with a broader epidermal pattern of stable accessibility. Chicken keratin-domain peaks showed lower variability than matched background peaks.

**(M)** Candidate CRE counts by species and keratin cluster. CREs were defined from ATAC-seeded intervals within α-keratin domains and organized by species and type-I/type-II cluster.

**(N)** Chromatin-state composition of candidate CREs across species. CREs were classified using promoter proximity and histone-mark annotations into promoter-like distal, promoter-proximal, active-enhancer-like, primed-enhancer-like, bivalent/mixed, repressed/polycomb-associated, or accessible-unclassified states. Differences across species are interpreted descriptively because available chromatin datasets differ in developmental stage and tissue context.

**(O)** ZEB2 motif enrichment at mouse CRE–KSR-loop-associated candidate CREs. HOMER known-motif analysis of mouse candidate CREs supported by both CRE–KSR contact and loop annotations. Bars show −log10 Benjamini q values for the ZEB2 motif across the CRE–KSR-loop foreground and comparison strata. Species-matched ATAC peaks excluding candidate CREs, candidate KSRs, and keratin promoter regions were used as background. Motif enrichment nominates ZEB2-associated regulatory input but does not establish ZEB2 occupancy or function at α-keratin loci.

**(P)** Exploratory motif enrichment at chicken bivalent/mixed candidate CREs. HOMER known-motif analysis of chicken candidate CREs classified as bivalent or mixed chromatin states. Bars show −log10 Benjamini q values for enriched RUNX1-, HOXA10-, and HOXD10-family motifs. The foreground contained 27 bivalent/mixed candidate CREs, and species-matched ATAC peaks excluding candidate CREs, candidate KSRs, and keratin promoter regions were used as background. Because of the limited foreground size and mixed chromatin-state classification, these results are interpreted as exploratory TF-family prioritization and do not establish TF occupancy, enhancer function, or direct regulation of keratin genes.

**(Q)** Conservative Tier-2 model. Within the KSR-defined scaffold, keratin promoters and distal candidate CREs occupy a broadly accessible, activity-marked regulatory landscape. These data support regulatory competence within keratin domains but do not define a universal enhancer–promoter code for individual keratin genes.

**Figure S4. Super-enhancer analysis and two-tier model of α-keratin cluster regulation.**

**(A)** FitHiC2 results showing interchromosomal contacts between the type-I keratin cluster on mouse chromosome 11 and the type-II keratin cluster on chromosome 15. Red lines indicate significant trans contacts. RNA-seq tracks show normalized read coverage in bins per million mapped reads. ChIP-seq tracks show fold enrichment over input. The mouse genome assembly used is GRCm38/mm10.

**(B)** Identification of clustered enhancers using ROSE. Enhancer regions were ranked to identify high-density enhancer clusters in skin or keratinocyte datasets.

**(C)** A/B compartment profiles at human α-keratin loci. Compartment correlation maps and first-eigenvector tracks for the human type-I locus on chr17 at D6 NHEK Hi-C. After PC1 sign orientation by active genomic features, type-I keratin loci are consistent with A-like compartment positioning, with the type-II locus appearing more boundary-proximal/mixed (data not shown).

**(D)** Chromosome overview of type-I and type-II α-keratin analysis windows in human, mouse, and chicken.

**(E)** Full-sample matched-null contact enrichment across human, mouse, and chicken. Per-sample rank percentile of the observed type-I × type-II α-keratin contact within the same five-covariate Mahalanobis matched-null framework for all 21 Hi-C/Micro-C samples. Samples are grouped by species and, for chicken, ordered by developmental and tissue context from embryonic dorsal epidermis to feather, scale, and brain controls. Open-ring markers indicate chicken brain controls, for which nearest-stage epidermal TMM was used as the expression covariate because matched brain RNA-seq was unavailable. This panel provides full-sample context for the replicate-resolved analysis in Figure 4D.

**(F)** Covariate-ablation sensitivity of the Mahalanobis matched-null test. Leave-one-out analysis for representative human D6, mouse P0 basal, and chicken E14 feather samples. Bars show the rank percentile of the observed type-I × type-II α-keratin contact under the full five-covariate matched-null model and after removing one covariate at a time: compartment PC1, gene density, state-matched TMM expression, GC content, or mappability proxy. The dotted line marks the 95th percentile.

**(G)** Predicted TF-binding motifs at ATAC-seq peaks within skin regional super-enhancers. Candidate TFs were prioritized based on motif presence together with RNA-seq expression or promoter-proximal accessibility, providing a complementary view of regulatory inputs outside the KSR-centered CRE catalog.

**Table S1. Comparative α-keratin gene registry across chicken, mouse, and human, related to Figure 1**. Curated registry of type-I and type-II α-keratin genes used for all cross-species analyses. The table includes build-matched genomic coordinates, gene identifiers, cluster assignments, reciprocal-hit orthology relationships, and synteny-based one-to-one or one-to-many assignments for chicken, mouse, and human α-keratin loci. This registry provides the reference gene set for comparative expression, chromatin-domain, candidate KSR, and candidate CRE analyses.

**Table S2. Dataset catalog for comparative α-keratin expression, chromatin, and 3D genome analyses, related to Figure 1**. Catalog of human, mouse, and chicken datasets used or evaluated in this study. For each dataset, the table lists species, genome build, assay type, sample identity, replicate group, tissue or cell source, developmental stage or cell state, anatomical region, accession or data source, and usage category. Assays include RNA-seq, ATAC-seq, histone-modification ChIP-seq, CTCF ChIP-seq or motif resources, DNA methylation, Hi-C, and Micro-C where available. This table documents sample provenance and build compatibility for the comparative multi-omic analyses.

**Table S3. Candidate Keratin Scaffold Regions across three species and two α-keratin clusters, related to Figure 2**. Genomic coordinates and feature annotations for the 36 active candidate Keratin Scaffold Regions (KSRs) identified in this study. Candidate KSRs are reported in native genome assemblies without liftOver: human GRCh38/hg38, mouse GRCm38/mm10, and chicken GRCg6a/galGal6. The active set includes 13 human KSRs, 17 mouse KSRs, and 6 chicken KSRs, distributed across type-I and type-II α-keratin clusters. For each candidate KSR, the table reports candidate identifier, species, genome assembly, α-keratin cluster, chromosome, 0-based half-open genomic coordinates, contact-boundary support, CTCF ChIP-seq overlap, CTCF motif support, ATAC accessibility, position class, distance to the nearest keratin-domain boundary, inclusion basis, and feature-interpretation notes. Position classes distinguish intra-cluster sub-boundaries, domain-flanking regions, and outer-flank sensitivity intervals. This table is the fixed denominator for Figure 2 and Figure S2 analyses; downstream KSR feature, motif, and control comparisons use this active set.

**Table S4. Candidate cis-regulatory elements and chromatin-state annotations across α-keratin loci, related to Figure 3**. Catalog of candidate cis-regulatory elements (CREs) within type-I and type-II α-keratin domains across human, mouse, and chicken. Candidate CREs were defined from species-matched ATAC-seq replicated or consensus peaks, merged within each species, and restricted to primary domain-bounded α-keratin analysis windows. The catalog contains 406 candidate CREs: 216 human, 130 mouse, and 60 chicken intervals, reported in native genome assemblies. For each CRE, the table reports genomic coordinates, species, cluster assignment, ATAC evidence across available states, accessibility state specificity, histone-mark availability and overlap, chromatin-state class, relationship to candidate KSRs, promoter proximity, and contact-support annotations. Chromatin-state classes were assigned using H3K27ac, H3K4me1, H3K4me3, H3K27me3, and canonical KRT transcription start-site proximity. Classes include promoter-proximal CRE, promoter-like distal, active-enhancer-like, primed-enhancer-like, bivalent or mixed, repressed or polycomb-associated, accessible-unclassified, and chromatin-annotation-unavailable. The table also includes CRE relationships to candidate KSRs and KRT promoters, including KSR overlap, distance to the nearest KSR, CRE–KSR contact support, CRE–promoter contact support, CRE–CRE contact support, TSS proximity layers, candidate target-gene annotations by proximity/contact/expression context, regulatory-mode interpretation, CTCF architectural advisory, claim level, and reviewer-risk flags. All CREs are candidate elements; the table does not imply validated enhancer activity, TF binding, conservation, or causal gene regulation.

**Table S5. Candidate CRE motif enrichment and TF-expression integration across three species, related to Figure 3 and Figure S3.** Consolidated motif-enrichment and transcription-factor expression table for candidate CRE strata across human, mouse, and chicken α-keratin loci. HOMER known-motif enrichment was performed across stratified subsets of the M03B candidate CRE catalog, using species-matched background intervals composed of pooled ATAC peaks with candidate CREs, active KSRs, and KRT promoter windows removed. Strata include species-level, regulatory-mode, KSR-proximity, and chromatin-state subsets. Benjamini q-values are reported as provided by HOMER. For each motif-stratum-species result, the table reports foreground definition, background definition, motif name, motif family, enrichment statistics, foreground size, power category, mapped transcription-factor gene, state-mean TF expression, and paralog-substitution status where applicable. Motif analyses are power-gated by foreground size: formal tests for n ≥ 20, exploratory tests for 10 ≤ n < 20, and presence-only summaries for n < 10. TF-expression integration was performed using full-transcriptome TMM matrices for each species. Expression values are summarized as state-mean log₂(TMM + 1) for the queried TF gene across available keratinocyte or epidermal states. The EKLF/KLF motif is mapped to KLF4 for keratinocyte interpretation, while KLF1 is retained only as the historical motif-training label. When canonical TF models were absent from a species expression matrix because of annotation gaps, the table records the substituted same-family paralog and flags the substitution. This applies to chicken JUNB-to-JUND and FOSL1-to-FOS substitutions. Motif results are used only for candidate prioritization and do not establish TF occupancy, conservation, enhancer function, or causal regulation.

**Table S6. Candidate inter-cluster contact-prioritized anchors between type-I and type-II α-keratin domains, related to Figure 4**. Candidate contact-prioritized anchors derived from the type-I × type-II α-keratin domain block, reported as a prioritization resource for targeted follow-up experiments (Capture-C, 4C, Oligopaint DNA-FISH, CRISPRi tiling) rather than as validated focal contacts. Each row is a candidate anchor: a bin-pair-level hot-spot within the type-I × type-II analysis window that exceeded the per-sample top-5% balanced contact threshold and merged by adjacency (within ± 1 bin in both axes) across at least one resolution among the four tested (64, 128, 256, 512 kb). For each candidate the table reports the type-I and type-II anchor coordinates on the species-native genome assembly (hg38, mm10, galGal6), the supporting state or replicate group, the number of biological replicates in which the anchor was recovered, the anchor’s overlap with candidate KSRs and with the candidate cis-regulatory element catalog, motif-integration evidence, the composite priority score, the recommended validation assay, and the candidate scope. Pixel-level inter-replicate reproducibility at current Hi-C / Micro-C depth is insufficient to elevate any single anchor from prioritization to focal mechanism (top-5% Jaccard within human replicate pairs is INSUFFICIENT_DEPTH; mouse P0 basal × suprabasal Jaccard is 0.0–0.125; only 1 of 59 anchor_class A replicated anchors is fully pixel-replicated at the resolutions tested); the table therefore reports all 93 candidates as supplementary prioritization material and the main-figure claim (Figure 4B–D) is restricted to the domain-level enrichment statistic.

**Video S1. Dynamic chromatin structures of avian α-keratin gene clusters during embryonic skin development, related to Figure 2**.

## STAR★METHODS

### KEY RESOURCES TABLE

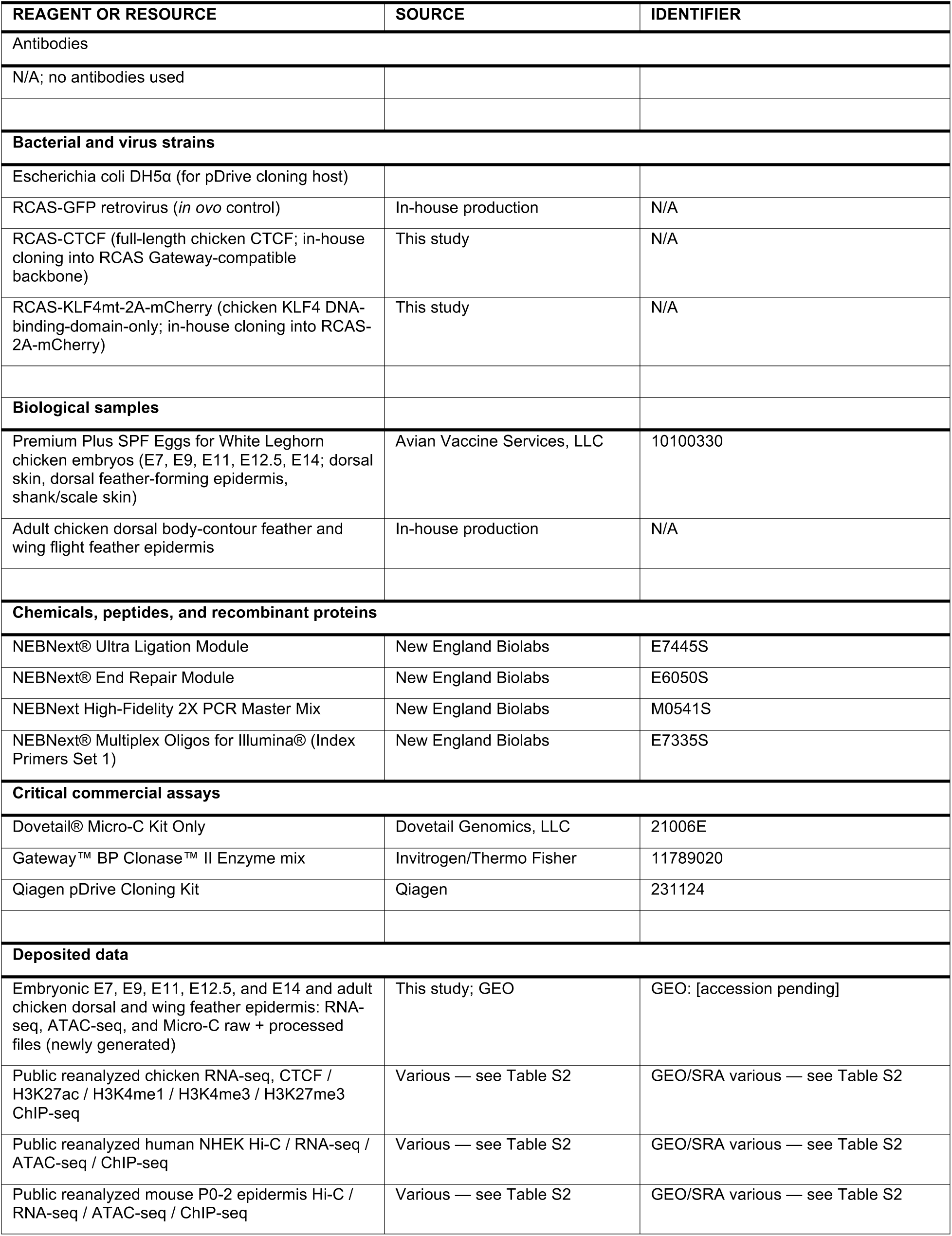
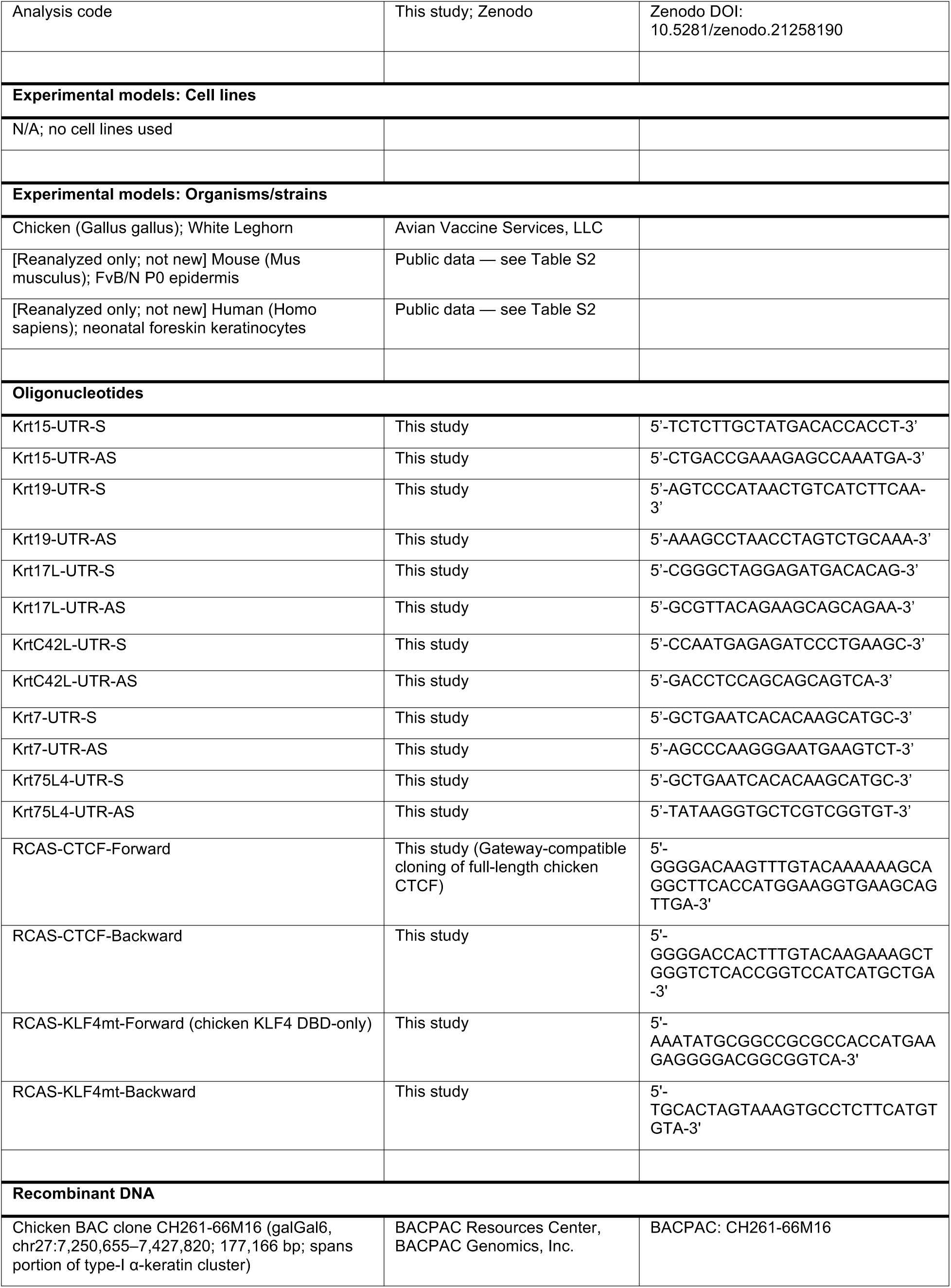
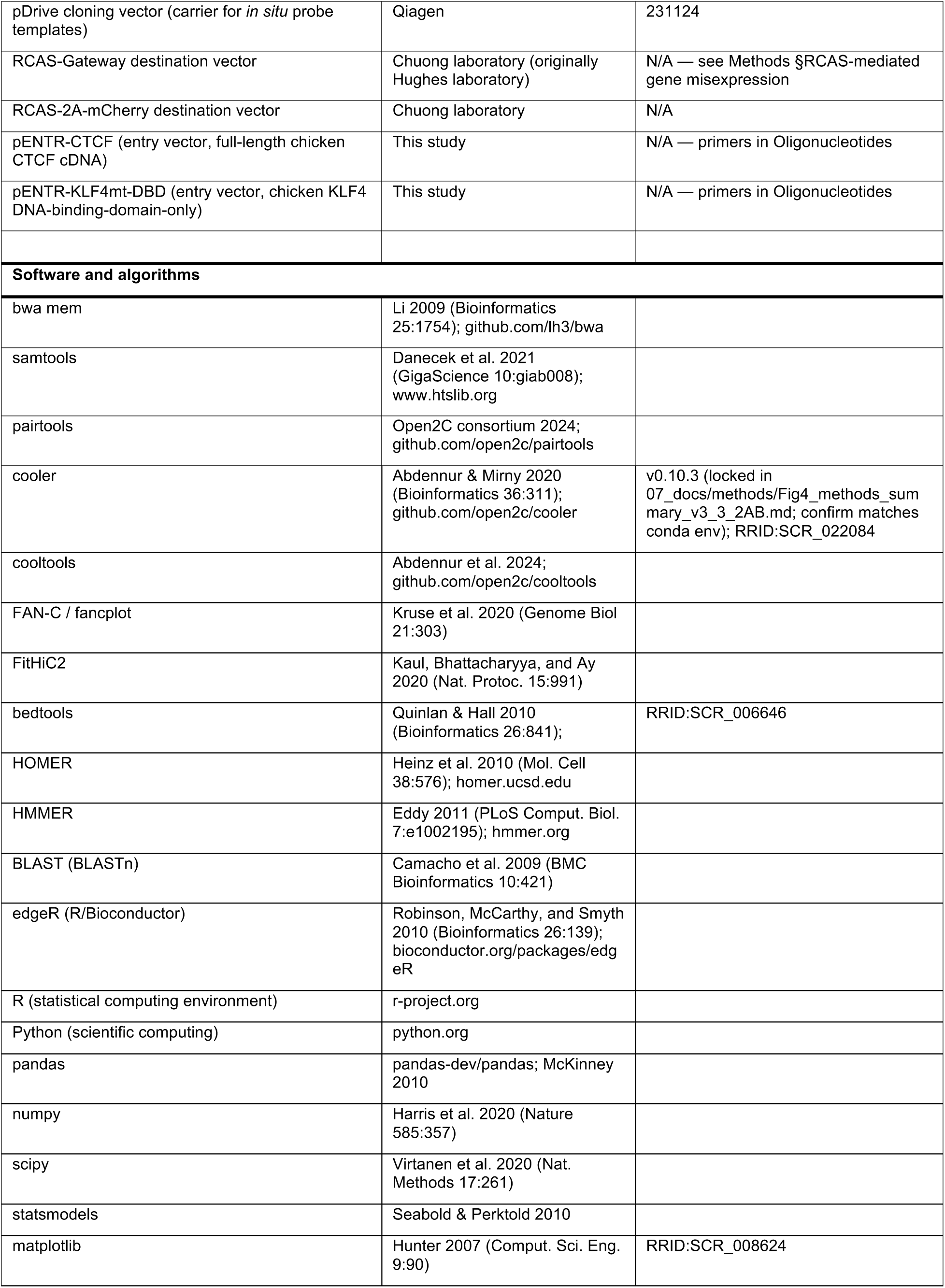
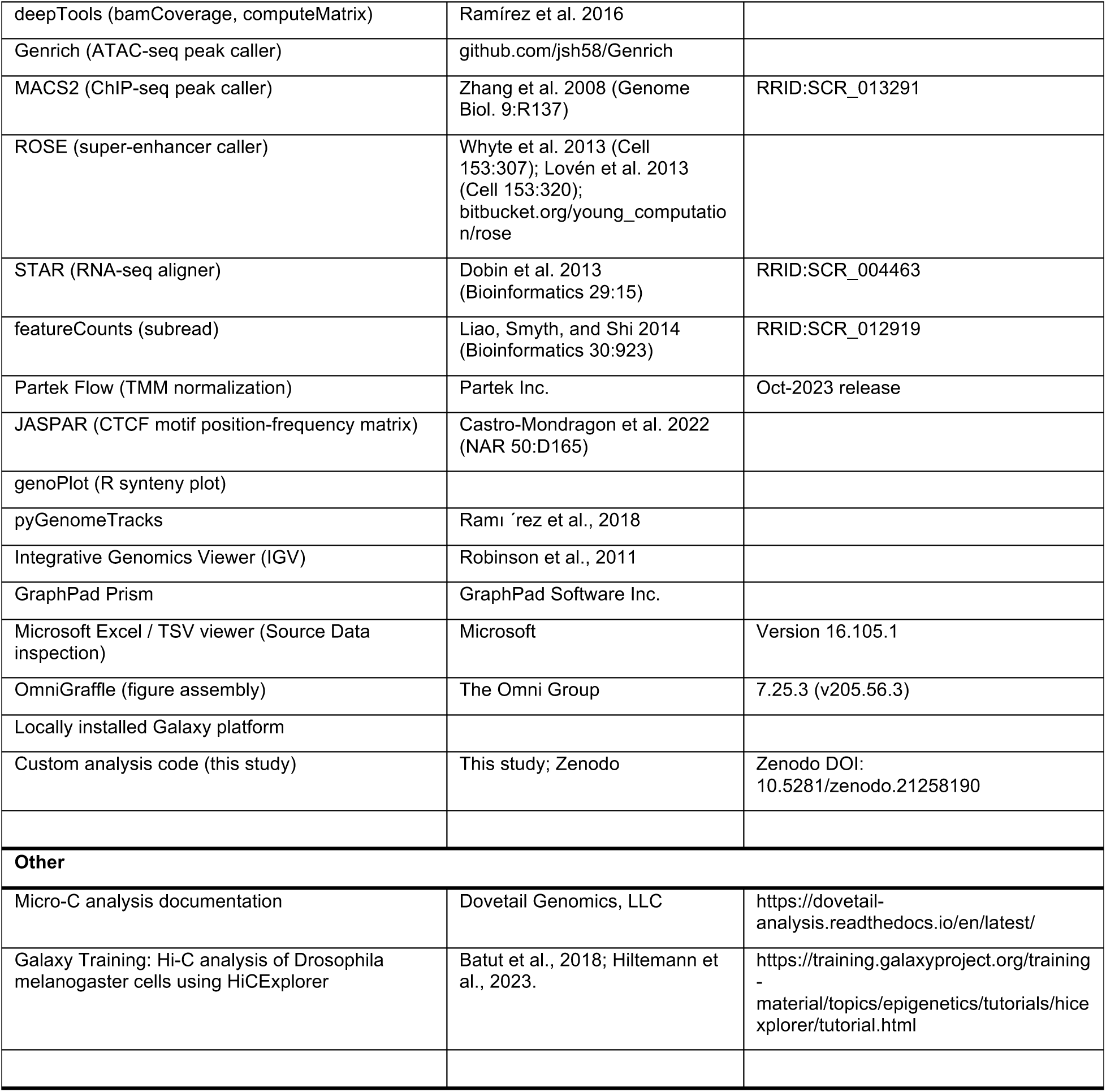

## EXPERIMENTAL MODEL AND STUDY PARTICIPANT DETAILS

### Chicken embryos and epidermal tissues

White Leghorn chicken embryos were used for developmental epidermal analyses, including RNA-seq, Micro-C, ATAC-seq, *in situ* hybridization, and RCAS perturbation experiments where indicated. Fertilized eggs were incubated under standard avian embryo conditions until tissue collection. Embryonic tissues were collected at the indicated developmental stages, including E7, E9, E11, E12.5, and E14, from dorsal skin, dorsal feather-forming epidermis, and shank or scale-forming skin. Adult feather epidermal samples were collected from dorsal body-contour feathers and wing flight feathers where specified. Animal use and embryo collection were performed in accordance with institutional guidelines under approved IACUC protocol #21512.

The ten chicken Micro-C libraries used in this study are listed as MiC-1 to MiC-10 in Table S2. All chicken Micro-C analyses, figure labels, and RNA-seq state-matched expression covariates used the curated tissue and developmental-stage assignments reported in Table S2.

Chicken embryonic samples were not sexed before analysis. Therefore, sex was not incorporated as a biological variable in chicken developmental comparisons. The primary chicken variables analyzed were developmental stage, anatomical region, appendage state, assay type, and keratin-cluster identity.

### Published mouse epidermal datasets

Published mouse epidermal datasets were reanalyzed to compare α-keratin expression, chromatin accessibility, histone-mark profiles, and chromatin architecture between basal and suprabasal epidermal compartments. These datasets included neonatal mouse epidermal or keratinocyte populations with RNA-seq, ATAC-seq, Hi-C, and/or ChIP-seq data as available. No new mouse experiments were performed for this study. Source datasets, accessions, sample identities, assay types, replicate groups, genome builds, and study usage are listed in Table S2.

Mouse analyses were interpreted as epidermal-state comparisons. Sex information was not consistently available in the source datasets and was not incorporated as an analytical variable.

### Published human keratinocyte and epidermal datasets

Human analyses used previously published or publicly available primary human keratinocyte and epidermal datasets, including neonatal normal human epidermal keratinocytes across calcium-induced differentiation and adult keratinocyte or epidermal datasets where available. No new human participants were recruited, and no new identifiable human specimens were collected for this study. Human datasets were analyzed as de-identified public or published data under the terms of the original studies and repositories. Dataset accessions, donor annotations where available, assay types, genome builds, and study usage are listed in Table S2.

Human donor sex was available for some datasets but was not balanced or consistently available across all datasets. Gender, ancestry, race, and ethnicity were not consistently reported and were not analyzed.

### Sex and gender as biological variables

Sex and gender were not primary analytical variables in this study. Chicken embryos were not sexed, mouse source datasets did not provide sex information in a form that could be consistently incorporated, and human donor metadata were incomplete across datasets. The major biological variables analyzed were species, developmental stage, anatomical region, epidermal differentiation state, assay type, and keratin-cluster identity. Therefore, this study was not designed to test sex- or gender-associated effects.

## METHOD DETAILS

### Synteny and α-keratin gene registry

α-Keratin gene annotations were curated for human, mouse, and chicken and assigned to type-I or type-II keratin clusters according to genomic position, gene identity, and syntenic context. Orthology relationships between human and mouse and between mouse and chicken were assigned from reciprocal ortholog hits. One-to-one orthologs were defined by unique reciprocal hits in pairwise BLASTn comparisons, whereas lineage-specific expansions were defined by one-to-many hits with similar alignment scores and sequence identities. Syntenic gene organization was visualized with the genoPlot package. Curated α-keratin gene identities, orthology assignments, cluster assignments, and synteny annotations are reported in Table S1.

### Chicken tissue collection, sectioning, and *in situ* hybridization

White Leghorn chicken embryos were collected at the indicated developmental stages and fixed in 4% paraformaldehyde at 4°C overnight. Tissues were dehydrated, embedded in paraffin, and sectioned at 7 μm thickness using a Leica microtome. *In situ* hybridization was performed as described previously^22^, with minor modifications. Gene-specific antisense RNA probes were generated from the 3′ untranslated regions of selected α-keratin transcripts to improve specificity among closely related keratin paralogs.

PCR products were cloned into the pDrive plasmid vector, and antisense RNA probes were synthesized from cloned inserts. Sections were faintly counterstained with diluted 2% eosin after hybridization. Images were acquired using a KEYENCE BZ-X710 microscope. Probe identities and primer sequences are provided in the Key Resources Table or Table S2.

### RNA-seq processing and TMM-normalized expression matrices

RNA-seq datasets from chicken epidermal tissues, human keratinocyte differentiation states, and mouse epidermal cell states were used to quantify gene expression across species-matched α-keratin loci. Sample identities, tissue or cell-state assignments, genome builds, and accession information are reported in Table S2.

Gene-level RNA-seq count matrices were processed in Partek Flow using a standard RNA-seq workflow and normalized by the trimmed mean of M values (TMM) method^38^. TMM normalization was performed separately within each species-specific dataset to account for library-size and composition differences. TMM-normalized matrices were used for keratin-gene expression summaries, transcription-factor expression annotation, and state-matched expression covariates in contact-enrichment analyses.

For expression heatmaps, TMM values were transformed as log₂(TMM + 1). Biological replicates were grouped by curated state labels, and state-mean expression was calculated for each gene. Heatmap values were gene-centered within each species by subtracting each gene’s mean across quantified states. No standard-deviation scaling was applied. Heatmap values therefore represent relative state bias for each gene, not absolute expression magnitude across genes or species.

Yanai τ^46^ specificity scores were calculated from linear TMM state means within each species and used as descriptive indices of expression breadth. The full α-keratin registry denominator was retained for class-count summaries, and quantified-gene counts were reported separately in the source data. Because the mouse dataset contains only basal and suprabasal states, mouse τ values were interpreted as basal– suprabasal bias rather than multi-state specificity.

For contact analyses, each Hi-C or Micro-C sample was paired with the closest RNA-seq state from the same species, tissue, developmental stage, or cell state using curated annotations in Table S2. State-matched TMM expression was summarized over genes overlapping each genomic window and used as one covariate in the five-dimensional Mahalanobis matched-null model.

### ATAC-seq library preparation and peak processing

ATAC-seq libraries generated in this study were prepared using an Omni-ATAC-based protocol. Viable cells or dissected epidermal samples were processed to recover nuclei under detergent-assisted lysis conditions. For sorted-cell preparations, approximately 50,000 cells were lysed in cold ATAC resuspension buffer containing NP-40, Tween-20, and digitonin. Nuclei were washed, transposed with Tn5 transposase at 37°C for 30 min, purified, pre-amplified, and subjected to qPCR to determine the required number of additional PCR cycles. Final libraries were purified, quantified, assessed for size distribution, and sequenced as paired-end libraries.

Chicken ATAC-seq reads were aligned to the galGal6/GRCg6a genome assembly using BWA. Aligned BAM files were name-sorted with samtools before peak calling. Peaks were called with Genrich using name-sorted BAM files as input, with narrowPeak and bedGraph signal outputs enabled. Genrich was run in ATAC-seq mode with mitochondrial reads excluded. Genrich narrowPeak files were used as the primary chicken ATAC peak calls. For visualization, Genrich-derived bedGraph files were cleaned, sorted, and converted to bigWig format with bedGraphToBigWig using the galGal6/GRCg6a chromosome-size file.

Mouse ATAC-seq data were processed in Partek Flow. Peaks were called with MACS2 v2.1.1 from paired-end BAM files in BAMPE mode using an effective genome size of 1.87 × 10⁹, q-value cutoff of 0.05, -- nomodel, --shift 37, --extsize 73, --call-summits, duplicate-retention threshold of 999, and bedGraph output enabled. Mouse ATAC-seq peaks were analyzed in mm10 coordinates.

Human neonatal keratinocyte ATAC-seq data were obtained directly from ENCODE for D0, D3, and D6 NHEK differentiation states. ENCODE fold-change-over-control bigWig signal tracks and replicated peak BED files were used directly and analyzed in hg38 coordinates.

For each species, state-level ATAC peak files were sorted and merged across matched epidermal or keratinocyte states to generate consensus accessible intervals. Consensus peaks were intersected with α-keratin domain windows and used for candidate CRE identification, candidate KSR annotation, chromatin-state classification, and regulatory-state comparisons.

### Published ChIP-seq datasets and reanalysis

All ChIP-seq datasets used in this study were previously published or publicly available. Chicken ChIP-seq data were reanalyzed using the previously described workflow^26^, including H3K27ac, H3K4me1, H3K4me3, CTCF, KLF-family, and SATB-family, with matching input or IgG controls where available. Human and mouse ChIP-seq datasets were obtained from published keratinocyte or epidermal resources listed in Table S2.

Chicken ChIP-seq data were reanalyzed using the workflow described in Liang et al. 2020. FASTQ files were groomed, quality-filtered, and quality-trimmed using a sliding window of 3 bases and a minimum quality score of 20. Reads were aligned with Bowtie2 using the --very-sensitive preset. Aligned reads were filtered with SAMtools using a minimum mapping quality of 1, sorted, and deduplicated.

Peaks were called with MACS2. For H3K27ac ChIP-seq, MACS2 was run with --nomodel, extension size 147 bp, bedGraph output enabled, and broad peak calling with broad-cutoff 0.1. For CTCF, KLF-family, and SATB-family ChIP-seq, MACS2 was run with --nomodel, bedGraph output enabled, and extension sizes estimated with MACS2 predictd. MACS2 bedGraph outputs were converted into fold-enrichment and log-likelihood-ratio signal tracks using macs2 bdgcmp, and signal tracks were converted to bigWig format for genome-browser visualization and locus-level plotting.

All ChIP-seq peaks and signal tracks were analyzed in the coordinate system used for the corresponding species-specific α-keratin figures: hg38 for human, mm10 for mouse, and galGal6/GRCg6a for chicken. When published files were generated in a different assembly, coordinates or signal tracks were harmonized before intersection with α-keratin features, as documented in Table S2.

ChIP-seq peaks and signal tracks were intersected with candidate KSRs, candidate CREs, keratin promoters, super-enhancer intervals, and contact-prioritized regions. H3K27ac and H3K4me1 were used to annotate enhancer-like chromatin; H3K4me3 and promoter proximity were used to annotate promoter-proximal or promoter-like chromatin; H3K27me3 was used to annotate polycomb-associated or repressed chromatin where available; and CTCF ChIP-seq was used as architectural annotation.

### Micro-C library preparation and processing

Chicken Micro-C libraries were generated using the Dovetail Micro-C Kit according to the manufacturer’s protocol, with sample-specific handling for embryonic and adult epidermal tissues. Dissected epidermal samples were crosslinked, subjected to *in situ* MNase digestion, lysed, captured on chromatin-capture beads, end-polished, bridge-ligated, proximity-ligated, reverse-crosslinked, and purified. Illumina-compatible libraries were prepared by end repair, adaptor ligation, USER digestion, streptavidin-based ligation-junction capture, index PCR, and size selection. Final libraries were quantified and assessed for size distribution before paired-end Illumina sequencing.

Micro-C reads were aligned to the chicken GRCg6a/galGal6 reference genome. The reference FASTA was indexed with samtools, a genome-size file was generated from the FASTA index, and a BWA index was built from the same reference. Paired-end reads were aligned with BWA-MEM using split-read-aware parameters. Aligned reads were parsed with pairtools using a minimum mapping quality of 40, sorted, deduplicated, and split into .pairs and BAM outputs. BAM files were coordinate-sorted and indexed with samtools. Pairtools duplicate statistics and preseq library-complexity estimates were retained for sample-level quality control.

Deduplicated valid-pair files were converted into Juicer-compatible .hic contact matrices using Juicer Tools. Contact maps were used for locus visualization, loop or focal-contact calling, TAD calling, A/B compartment analysis, and type-I × type-II keratin-domain contact analyses. Candidate loops or focal contacts were called with Mustache where applicable and treated as computational annotations. TADs were called from .hic matrices using Juicer Tools Arrowhead with Knight–Ruiz (KR) normalization where appropriate. Additional contact-map visualization and matrix-based analyses followed HiCExplorer-style workflows.

A/B compartment profiles were computed from processed contact matrices using FAN-C^47^. Compartment correlation matrices were generated with fanc compartments, first-eigenvector tracks were exported with the -v option, and compartment maps were visualized with fancplot. Eigenvector sign was interpreted relative to active genomic features so that positive values corresponded to A-like chromatin and negative values corresponded to B-like chromatin. Compartment PC1 values were also used as a covariate in the Mahalanobis matched-null contact-enrichment analysis.

### Human and mouse Hi-C reprocessing and analysis

Published human keratinocyte and mouse epidermal Hi-C datasets were reprocessed or analyzed to generate contact maps and chromatin-architecture annotations at α-keratin loci. Human NHEK reads were aligned to GRCh38/hg38, and mouse epidermal reads were aligned to GRCm38/mm10. Sample identities, accessions, genome builds, and processing status are reported in Table S2.

Mouse paired-end Hi-C reads were aligned with BWA-MEM, parsed with pairtools using a minimum mapping quality of 40, sorted, deduplicated, and exported as .pairs and BAM files. Deduplicated .pairs files were converted into Juicer-compatible .hic contact maps and cooler-compatible .cool/.mcool matrices^48^. Multi-resolution balanced matrices were generated with cooler for downstream visualization and quantitative analyses.

Human NHEK datasets were analyzed within the same downstream contact-map framework after BWA-MEM alignment and generation of Juicer- and cooler-compatible matrices where available. Because the retained human command record did not preserve the complete pairtools command history, human and mouse datasets were treated as jointly analyzed contact-map resources rather than as datasets generated by identical command lines.

Candidate loops or focal contacts were called from .hic contact maps using Mustache on the α-keratin chromosomes: human chr17 and chr12, and mouse chr11 and chr15. TADs were called using Juicer Tools Arrowhead at 10-kb and/or 25-kb resolution where sufficient data were available. BAM-derived coverage tracks were generated with deepTools bamCoverage for visualization and quality-control support. FAN-C was used for A/B compartment profiles. Loop, TAD, boundary, and compartment features were treated as computational annotations of chromatin architecture.

### Identification of candidate Keratin Scaffold Regions

Candidate Keratin Scaffold Regions were identified within species- and cluster-specific α-keratin-domain discovery windows. These windows were defined from the curated α-keratin gene registry together with contact-derived domain structure at each type-I and type-II locus. Final discovery-window coordinates and candidate KSR coordinates are reported in the Figure 2 source data and Table S3.

Contact-derived boundary evidence was the primary criterion for KSR discovery. Boundary positions were obtained from hicFindTADs calls on normalized Hi-C or Micro-C contact matrices binned at 10-kb resolution. Boundaries were defined from local minima of the multiscale TAD-insulation score. ATAC-seq, histone modifications, RNA-seq, CTCF ChIP-seq, and motif data were not used to call boundaries; these datasets were added after boundary discovery as annotation layers.

For each species and α-keratin cluster, contact-derived boundary records were restricted to the final discovery window, sorted by coordinate, and merged when adjacent records were separated by no more than 25 kb. Each merged interval was treated as a consensus boundary seed. Supporting contact datasets, developmental or cell-state groups, insulation scores, and reproducibility classes were recorded for each seed.

Consensus boundary seeds were extended 15 kb upstream and downstream to generate candidate KSR intervals. Overlapping extended intervals were merged and clipped to the final discovery window. Candidate intervals were assigned position classes relative to the keratin cluster. The active candidate KSR set used for downstream analyses contained 36 intervals across the three species: 13 human, 17 mouse, and 6 chicken KSRs.

After candidate intervals were defined, each KSR was annotated with CTCF ChIP-seq signal, CTCF motif evidence, ATAC-seq accessibility, histone-mark overlap, loop or contact-anchor support, and nearby keratin-gene expression. Annotation layers did not alter KSR positions or determine whether a candidate was initially called.

Active candidate KSRs were compared with matched control intervals, including within-TAD random intervals, CTCF-anchor-matched intervals, loop-anchor-matched intervals, non-keratin gene-cluster boundaries, and expressed-gene boundaries outside keratin clusters. Foreground and control intervals were scored with the same feature pipeline, including TAD-boundary overlap, CTCF ChIP-seq peak count, CTCF motif count, ATAC peak count, histone-mark overlap, loop-anchor count, and mean TAD-separation score.

CTCF motif orientation was analyzed as an architectural annotation. Strand-aware CTCF motif calls were intersected with CTCF ChIP-seq peaks and candidate KSR intervals. Motifs were classified as convergent, divergent, tandem, single motif, mixed or multiple, or no motif. Orientation classes were compared with matched control intervals using the same motif source and classification rules.

Accessible sequences within candidate KSRs were analyzed for transcription-factor motif enrichment using HOMER known-motif analysis. ATAC-seq peaks overlapping active candidate KSRs were used as foreground intervals, and species-matched genome-wide ATAC peaks were used as background. Motif enrichment was interpreted as sequence-based prioritization and not as evidence of direct TF occupancy or regulatory function.

### Identification of candidate cis-regulatory elements

Candidate cis-regulatory elements were defined from species-matched ATAC-seq peaks within α-keratin domain windows. The domain-bounded windows were defined from the curated α-keratin registry and boundary-bounded keratin-domain framework and analyzed in native genome assemblies without liftOver. Window coordinates are reported in the Figure 3 source data.

Human CREs were seeded from replicated ENCODE ATAC-seq peaks from D0, D3, and D6 neonatal keratinocytes. Mouse CREs were seeded from MACS2 narrowPeak files from P0 basal and suprabasal epidermal samples. Chicken CREs were seeded from ATAC-seq narrowPeak files from E7 dorsal epidermis, E9 dorsal epidermis, E14 dorsal feather epidermis, and E14 leg/scale epidermis.

For each species, per-state ATAC peaks were sorted and merged across states using zero-gap merging to generate a state-pooled consensus accessible-peak set. Consensus peaks were intersected with type-I or type-II keratin-locus analysis windows, and intervals outside the domain-bounded keratin windows were excluded. This procedure generated 406 candidate CREs: 216 human, 130 mouse, and 60 chicken CREs.

Candidate KSRs were used as annotation features, not as CRE discovery seeds. Each CRE was annotated for KSR overlap, promoter proximity, chromatin-contact support, chromatin-state class, H3K27ac status, and nearest keratin gene. Promoter-proximal CREs were defined by overlap with canonical keratin transcription start-site windows within ±2 kb. Contact-support categories were assigned using species-matched Hi-C or Micro-C loop/contact calls and included CRE–KSR, CRE–promoter, and CRE–CRE support.

Candidate CREs were classified using available ATAC-seq and ChIP-seq features, including H3K27ac, H3K4me1, H3K4me3, H3K27me3, and promoter proximity. CREs were assigned to promoter-proximal, promoter-like distal, active-enhancer-like, primed-enhancer-like, bivalent or mixed, repressed/polycomb-associated, accessible-unclassified, or annotation-unavailable classes.

To summarize accessibility and activity-associated chromatin behavior, CREs were also assigned to collapsed ATAC–H3K27ac classes. ATAC accessibility was considered stable when the maximum absolute pairwise log₂ fold-change across matched states was <0.5 and dynamic when it was ≥0.5. H3K27ac dynamics were classified according to the stage structure and data availability for each species. These classes were used for visualization and descriptive classification, not as formal differential-accessibility or enhancer-activity tests.

Motif enrichment was performed on stratified candidate CRE foregrounds using HOMER known-motif results. The primary background was the species-matched genome-wide ATAC peak set after excluding candidate CREs, active candidate KSRs, and keratin promoter windows. Motif enrichments were paired with state-mean transcription-factor expression from TMM-normalized RNA-seq matrices to prioritize candidate TF families. Motif results were not interpreted as evidence of TF occupancy or regulatory function.

### Interchromosomal contact prioritization using FitHiC2

Interchromosomal type-I × type-II α-keratin contact candidates were prioritized using FitHiC2^39^ on processed Hi-C or Micro-C contact data. Candidate contacts were analyzed between species-matched type-I and type-II α-keratin chromosomes: human chr17 × chr12, mouse chr11 × chr15, and chicken chr27 × chr33.

For each sample and resolution, contact-count files and fixed-bin fragment files were generated. Bin-level bias files were generated with HiCKRy and supplied to FitHiC2. FitHiC2 was run in interchromosomal mode using the interOnly option. Multiple resolutions were analyzed where sufficient data were available. FitHiC2 outputs were filtered to retain contacts overlapping the type-I × type-II α-keratin analysis-window block.

Candidate contact bins were intersected with keratin-domain annotations, including candidate KSRs, TAD or sub-TAD boundaries, candidate CREs, keratin promoters, and super-enhancer intervals. FitHiC2 was used as a contact-prioritization tool. Enriched interchromosomal bin pairs were interpreted as population-level contact enrichment and not as validated single-cell pairing, focal physical anchors, or causal regulatory loops.

### Super-enhancer identification using ROSE

Super-enhancer annotations were generated from activity-associated chromatin intervals using ROSE. Candidate enhancer input regions were derived from H3K27ac-enriched and H3K4me1-enriched peak sets. Peak files were annotated with HOMER, converted to BED/GFF-compatible formats, and supplied to ROSE with species-matched genome annotations.

Enhancer regions were stitched using the ROSE default 12-kb stitching distance. Regions within transcription start-site exclusion windows were removed according to the ROSE workflow. H3K27ac ChIP-seq BAM files and matched input-control BAM files were mapped to stitched enhancer regions. Input-subtracted H3K27ac signal was calculated for each stitched enhancer, with negative values set to zero before ranking.

Stitched enhancers were ranked by input-subtracted H3K27ac signal. ROSE identified the super-enhancer cutoff using the tangent-point method on the ranked enhancer distribution. Regions above this cutoff were classified as super-enhancers, and remaining stitched regions were classified as typical enhancers. ROSE-defined super-enhancers were intersected with α-keratin-domain windows, candidate KSRs, candidate CREs, promoters, and FitHiC2-prioritized contact regions.

### Type-I × type-II α-keratin matched-null contact enrichment

Type-I × type-II α-keratin interchromosomal contact enrichment was quantified for each Hi-C or Micro-C sample. For each species, type-I and type-II analysis windows were defined from the curated α-keratin registry by spanning the corresponding keratin cluster and flanking the locus to capture the local keratin-domain context. Exact window coordinates are reported in the Figure 4 source data.

For each sample, the observed type-I × type-II contact statistic was calculated at 128-kb resolution from the rectangular contact submatrix spanning the species-matched type-I and type-II analysis windows. The observed value was defined as the mean iterative correction and eigenvector decomposition (ICE)-balanced^49^ contact across non-zero bin pairs for which both bins had finite balancing weights. The same filtering rule was applied to observed keratin-domain pairs and matched non-keratin background pairs.

To test whether the observed contact exceeded expectation for comparable non-keratin regions, a sample-specific matched-null distribution was constructed using five-dimensional Mahalanobis nearest-neighbor matching in covariate space^40^. Matching was performed at 256-kb resolution using five window-level covariates: A/B-compartment PC1, gene density, state-matched TMM expression, GC fraction, and a matrix-visibility/mappability proxy. The mappability proxy was derived from cooler bin weights after matrix balancing^48^ and was not treated as equivalent to sequence-level UMAP mappability.

Candidate background windows were drawn from the same chromosomes as the corresponding type-I and type-II keratin windows, excluding keratin-domain windows. For each chromosome, the 200 non-keratin windows nearest to the keratin-window covariate profile were selected using Mahalanobis distance computed from the candidate-pool covariance matrix with 10⁻⁶ ridge regularization. Pairing the 200 matched windows from the type-I chromosome with the 200 matched windows from the type-II chromosome generated 40,000 matched non-keratin window pairs per sample.

For each matched non-keratin pair, the mean 128-kb ICE-balanced contact value was calculated from the corresponding rectangular submatrix using the same non-zero and finite-weight filtering applied to the observed keratin block. Empirical *p*-values were calculated as the fraction of matched-null pairs with mean balanced contact greater than or equal to the observed type-I × type-II keratin contact. Rank percentile was calculated as the fraction of matched-null pairs with mean balanced contact lower than the observed value, multiplied by 100. A 95% confidence interval for the rank percentile was estimated by 1,000 bootstrap resamplings of the matched-null pool with replacement.

Figure 4D reports the replicate-resolved analysis across seven biological-state replicate pairs: human D0, D3, and D6 keratinocytes; mouse P0 basal and suprabasal epidermis; chicken E14 dorsal feather epidermis; and chicken E14 leg scale epidermis. Figure S4E reports the same statistic across the full 21-sample Hi-C/Micro-C set, including chicken brain controls. Because matched chicken brain RNA-seq was unavailable, brain-control matched-null tests used the nearest-stage epidermal TMM profile as the expression covariate.

Leave-one-out covariate-ablation analysis was performed for representative human D6, mouse P0 basal, and chicken E14 feather samples by repeating Mahalanobis matching after removing one covariate at a time. The resulting rank percentiles were compared with the full five-covariate model and are shown in Figure S4F.

### RCAS-mediated gene misexpression

For RCAS-mediated CTCF misexpression, full-length chicken CTCF was cloned into RCAS using the Gateway system. Chicken E14 dorsal skin cDNA was used as PCR template. For RCAS-mediated expression of a dominant-negative KLF4 construct, the DNA-binding domain was amplified and ligated into an RCAS vector carrying an mCherry fluorescent tracer. RCAS-GFP virus was used as control. Primer sequences are reported in the Key Resources Table or Table S2.

RCAS viruses were prepared and concentrated by ultracentrifugation at 20,000 rpm for 2 h at 4°C. Virus was injected into the amniotic cavity at E3 at 1 × 10⁷ IU/ml, and embryos were collected between E13 and E15 for downstream analysis.

### Candidate-feature interpretation

Candidate KSRs, candidate CREs, FitHiC2-prioritized contacts, contact-prioritized regions, and ROSE-defined super-enhancers were treated as computationally nominated features unless directly tested by perturbation or another orthogonal assay. Contact-map enrichments from Hi-C and Micro-C were interpreted as population-level contact signals and not as single-cell pairing, deterministic physical co-localization, or validated causal enhancer–promoter loops.

## QUANTIFICATION AND STATISTICAL ANALYSIS

### General statistical framework

Statistical analyses were performed within species unless otherwise indicated. No direct cross-species statistical comparison of absolute expression, chromatin signal, or contact magnitude was performed. Human, mouse, and chicken analyses were compared at the level of qualitative pattern, matched feature classes, and species-specific statistical support. Source data report sample identifiers, biological-state assignments, genome builds, replicate structure, statistical tests, effect sizes, *p*-values, adjusted *p*-values, and analysis-specific denominators.

### α-Keratin expression state bias

For Figure 1, biological replicates were grouped by curated epidermal state, and state-mean expression was calculated from log₂(TMM + 1) values. Heatmap values were centered within each gene and species by subtracting the mean across quantified states. No standard-deviation scaling was applied.

Yanai τ specificity scores were calculated from linear state-mean TMM values and used as descriptive expression-breadth indices. Genes were assigned to predefined expression-breadth classes. These classes were not treated as formal differential-expression calls. Genes with missing, one-state-only, or zero-variance expression patterns were flagged in the source data.

### KSR matched-control and CTCF-orientation analyses

For candidate KSR matched-control analyses, active KSRs were compared with matched control intervals using the same feature-scoring pipeline. Statistical comparisons were performed for each species, feature, and control set using Mann–Whitney U tests and paired Wilcoxon signed-rank tests where appropriate. Cliff’s δ was calculated as an effect-size measure. Benjamini–Hochberg correction was applied within each test family. A comparison was considered supported when the Mann–Whitney U Benjamini–Hochberg FDR was <0.1 and the absolute Cliff’s δ was ≥0.2.

CTCF motif-orientation enrichment was tested by comparing convergent versus non-convergent orientation between KSRs and matched control intervals using two-sided Fisher’s exact tests, followed by Benjamini– Hochberg correction.

### CRE accessibility, chromatin-state, and motif analyses

For candidate CRE analyses, ATAC-seq and H3K27ac signals were summarized within each species and state. ATAC accessibility was classified as stable when the maximum absolute pairwise log₂ fold-change across matched states was <0.5 and dynamic when it was ≥0.5. H3K27ac dynamics were classified according to data availability and stage structure for each species. The resulting ATAC–H3K27ac classes were used for descriptive visualization and were not formal differential-accessibility or enhancer-activity calls.

Motif enrichment at candidate CREs or KSR-associated accessible intervals was assessed with HOMER known-motif analysis^50^ using species-matched ATAC background sets. HOMER Benjamini q values were used as reported. Motif enrichments were interpreted as candidate TF-family prioritization and not as direct evidence of TF binding or function.

### FitHiC2 contact prioritization

FitHiC2 was used to prioritize enriched interchromosomal contacts between type-I and type-II α-keratin chromosomes. Resolution-specific contact files, fragment files, and HiCKRy-derived bias files were used as inputs. Significant or prioritized contacts were filtered to the keratin analysis windows and intersected with KSR, CRE, promoter, TAD-boundary, and super-enhancer annotations. FitHiC2 outputs were treated as candidate contact-prioritization results, not as evidence of validated physical co-localization.

### Matched-null type-I × type-II contact enrichment

For each sample, the observed 128-kb type-I × type-II keratin contact statistic was compared with 40,000 matched non-keratin window pairs selected by five-dimensional Mahalanobis nearest-neighbor matching. Empirical *p*-values were calculated as the fraction of matched-null pairs with contact values greater than or equal to the observed keratin-domain value. Rank percentile was calculated as the fraction of matched-null pairs below the observed value multiplied by 100. Bootstrap 95% confidence intervals for rank percentile were estimated from 1,000 resamplings of the matched-null pool.

The replicate-resolved analysis is reported in Figure 4D, the full 21-sample analysis in Figure S4E, and the leave-one-out covariate-ablation analysis in Figure S4F. Pearson correlation was used to compare summed keratin log₂(TMM + 1) expression with type-I × type-II contact z-score across matched states where reported. Small-n correlations were treated as descriptive.

### Reporting and reproducibility

All figure-level quantitative outputs are reported in the corresponding source-data files. Sample identities, accessions, genome builds, coordinate status, replicate labels, software versions, and major analysis inputs are reported in Table S2, figure source data, and deposited code. Interval operations were performed with BEDTools-compatible workflows and custom Python/R scripts. Statistical analyses used Python and R packages, including NumPy, pandas, SciPy, statsmodels, and standard R statistical functions.

## Notes

### Competing Interest Statement

The authors have declared no competing interest.

