## Supplementary Figure S1-S4 for "Hierarchical Gene Cluster Regulation Across Vertebrate Skins: Developmental Control of Keratin Gene Expression"

| | Species | Chr | Genomic Location | Size (bp) | Krt Gene # | Gene Names (in gene order) |
|---|---|---|---|---|---|---|
| Type-I α-Keratin Gene Cluster (acidic keratins) | Chicken | 27 | 27:7320674-7449377 | 128,703 | 14 | *Krt222, Krt12, Krt20, Krt23, Krt15, Krt19, Krt42L, Krt10, Krt9, Krt17L, Krtc42L, Krt42L_2, Krt13, Krt17* |
| | Mouse | 11 | 11:99232760-100269951 | 1,037,190 | 27 | *Krt222, Krt24, Krt25, Krt26, Krt27, Krt28, Krt10, Krt12, Krt20, Krt23, Krt39, Krt40, (Krtap genes), Krt33a, Krt33b, Krt34, Krt31, Krt32, Krt35, Krt36, Krt13, Krt15, Krt19, Krt9, Krt14, Krt16, Krt17, Krt42* |
| | Human | 17 | 17:40654665-41624575 | 969,911 | 28 | *KRT222, KRT24, KRT25, KRT26, KRT27, KRT28, KRT10, KRT12, KRT20, KRT23, KRT39, KRT40, (KRTAP genes), KRT33A, KRT33B, KRT34, KRT31, KRT37, KRT38, KRT32, KRT35, KRT36, KRT13, KRT15, KRT19, KRT9, KRT14, KRT16, KRT17* |
| Type-II α-Keratin Gene Cluster (basic/neutral keratins) | Chicken | 33 | 33:5187366-5408440 | 221,074 | 17 | *Krt80, Krt7, Krt84, Krt75, Krt6a, Krt5, Krt75L3, Krt75L4, Krt73, Krt1, Krt79L, Krt75L2, Krt71, Krt75L1, Krt4L, Krt8, Krt18* |
| | Mouse | 15 | 15:101348314-102032026 | 683,711 | 27 | *Krt80, Krt7, Krt87, Krt88, Krt81, Krt86, Krt83, Krt84, Krt82, Krt80, Krt75, Krt6b, Krt6a, Krt5, Krt71, Krt74, Krt72, Krt73, Krt2, Krt1, Krt77, Krt76, Krt4, Krt79, Krt78, Krt8, Krt18* |
| | Human | 12 | 12:52168996-52952906 | 783,910 | 26 | *KRT80, KRT7, KRT81, KRT83, KRT85, KRT84, KRT82, KRT75, KRT6B, KRT6C, KRT6A, KRT5, KRT71, KRT74, KRT72, KRT73, KRT2, KRT1, KRT77, KRT76, KRT3, KRT4, KRT79, KRT78, KRT8, KRT18* |

*genome versions: chicken, GRCg6a/gg6; mouse, GRCm38/mm10; human, GRCh38/hg38

Figure S2

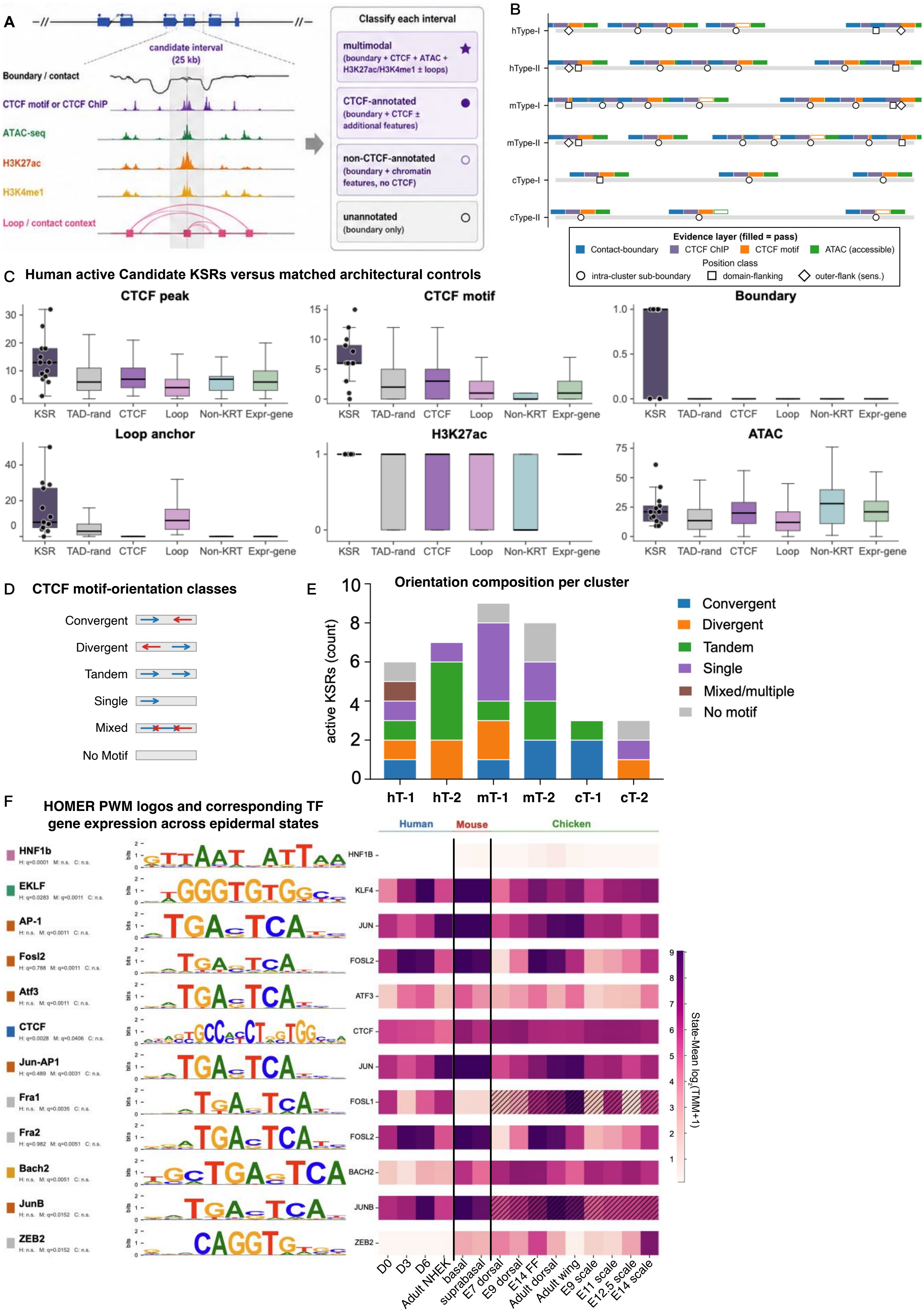
A
candidate interval (25 kb)
Boundary / contact
CTCF motif or CTCF ChIP
ATAC-seq
H3K27ac
H3K4me1
Loop / contact context
Classify each interval
multimodal (boundary + CTCF + ATAC + H3K27ac/H3K4me1 ± loops)
CTCF-annotated (boundary + CTCF ± additional features)
non-CTCF-annotated (boundary + chromatin features, no CTCF)
unannotated (boundary only)
B
hType-I
hType-II
mType-I
mType-II
cType-I
cType-II
Evidence layer (filled = pass)
Contact-boundary
CTCF ChIP
CTCF motif
ATAC (accessible)
Position class
intra-cluster sub-boundary
domain-flanking
outer-flank (sens.)
C
Human active Candidate KSRs versus matched architectural controls
CTCF peak
CTCF motif
Boundary
Loop anchor
H3K27ac
ATAC
KSR
TAD-rand
CTCF
Loop
Non-KRT
Expr-gene
D
CTCF motif-orientation classes
Convergent
Divergent
Tandem
Single
Mixed
No Motif
E
Orientation composition per cluster
active KSRs (count)
hT-1
hT-2
mT-1
mT-2
cT-1
cT-2
Convergent
Divergent
Tandem
Single
Mixed/multiple
No motif
F
HOMER PWM logos and corresponding TF gene expression across epidermal states
HNF1b
H: q=0.0001 M: n.s. C: n.s.
EKLF
H: q=0.0283 M: q=0.0011 C: n.s.
AP-1
H: n.s. M: q=0.0011 C: n.s.
Fosl2
H: q=0.788 M: q=0.0011 C: n.s.
Atf3
H: n.s. M: q=0.0011 C: n.s.
CTCF
H: q=0.0028 M: q=0.0406 C: n.s.
Jun-AP1
H: q=0.489 M: q=0.0031 C: n.s.
Fra1
H: n.s. M: q=0.0035 C: n.s.
Fra2
H: q=0.982 M: q=0.0051 C: n.s.
Bach2
H: n.s. M: q=0.0051 C: n.s.
JunB
H: n.s. M: q=0.0152 C: n.s.
ZEB2
H: n.s. M: q=0.0152 C: n.s.
Human
Mouse
Chicken
HNF1B
KLF4
JUN
FOSL2
ATF3
CTCF
JUN
FOSL1
FOSL2
BACH2
JUNB
ZEB2
D0
D3
D6
Adult NHEK
basal
suprabasal
E7 dorsal
E9 dorsal
E14 FF
Adult dorsal
Adult wing
E9 scale
E11 scale
E12.5 scale
E14 scale
State-Mean log2(TMM+1)

Figure S3 (A-C)

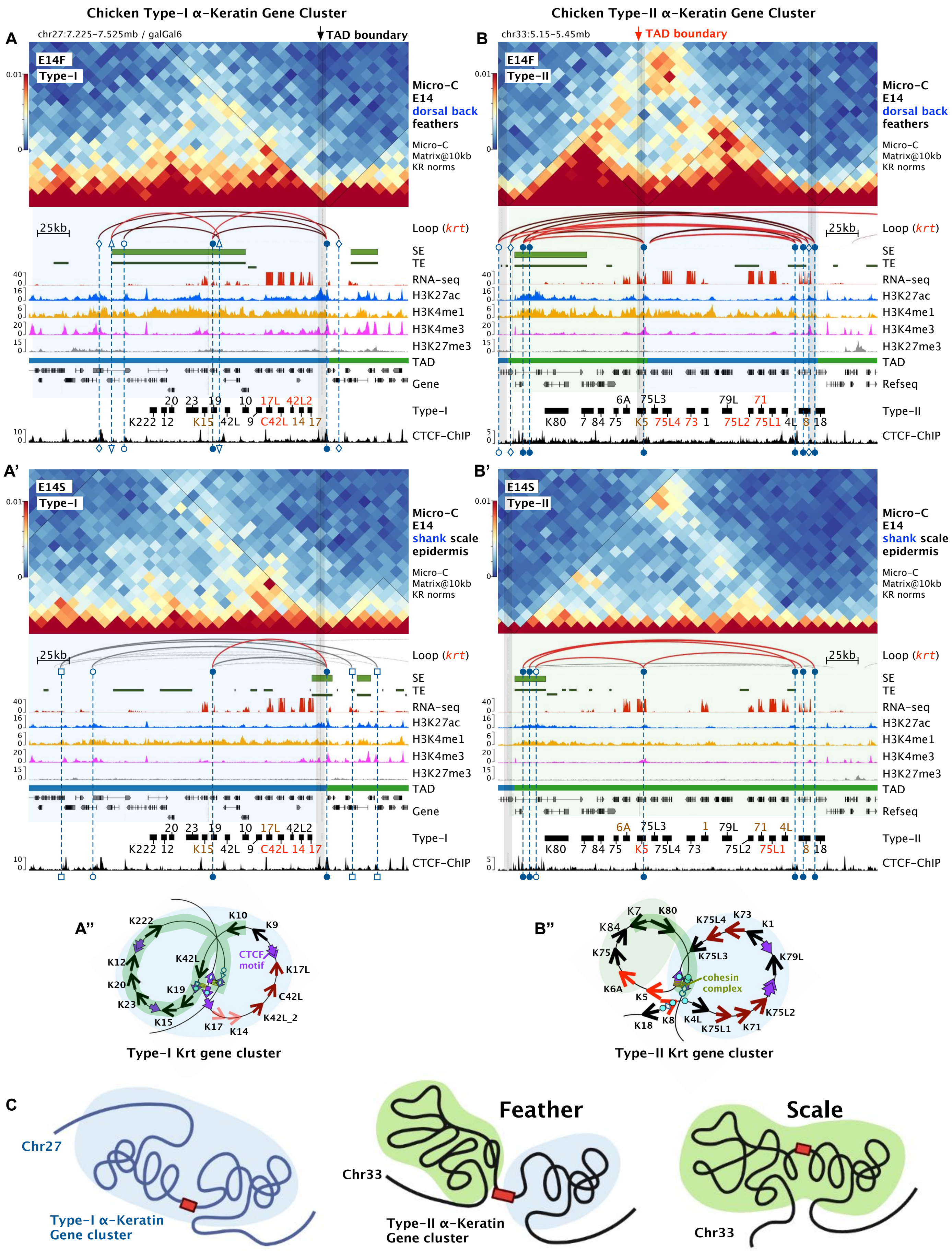

Chicken Type-I α-Keratin Gene Cluster
Chicken Type-II α-Keratin Gene Cluster
A
chr27:7.225-7.525mb / galGal6
TAD boundary
B
chr33:5.15-5.45mb
TAD boundary
E14F
Type-I
Type-II
Micro-C
E14
dorsal back
feathers
Micro-C
Matrix@10kb
KR norms
25kb
Loop (krt)
SE
TE
RNA-seq
H3K27ac
H3K4me1
H3K4me3
H3K27me3
TAD
Gene
Refseq
Type-I
Type-II
CTCF-ChIP
20 23 19 10 17L 42L2
K222 12 K15 42L 9 C42L 14 17
6A 75L3 79L 71
K80 7 84 75 K5 75L4 73 1 75L2 75L1 4L 8 18
A'
B'
E14S
shank scale
epidermis
6A 75L3 1 79L 71 4L
K80 7 84 75 K5 75L4 73 75L2 75L1 8 18
A"
B"
K222 K10 K9 K12 K42L CTCF motif K17L K20 K19 K23 C42L K15 K42L_2 K17 K14
Type-I Krt gene cluster
K7 K80 K84 K75L4 K73 K1 K75 K75L3 K79L K6A cohesin complex K5 K75L2 K18 K8 K4L K75L1 K71
Type-II Krt gene cluster
C
Chr27
Type-I α-Keratin
Gene cluster
Feather
Chr33
Type-II α-Keratin
Gene cluster
Scale
Chr33

Figure S3 (D-E)

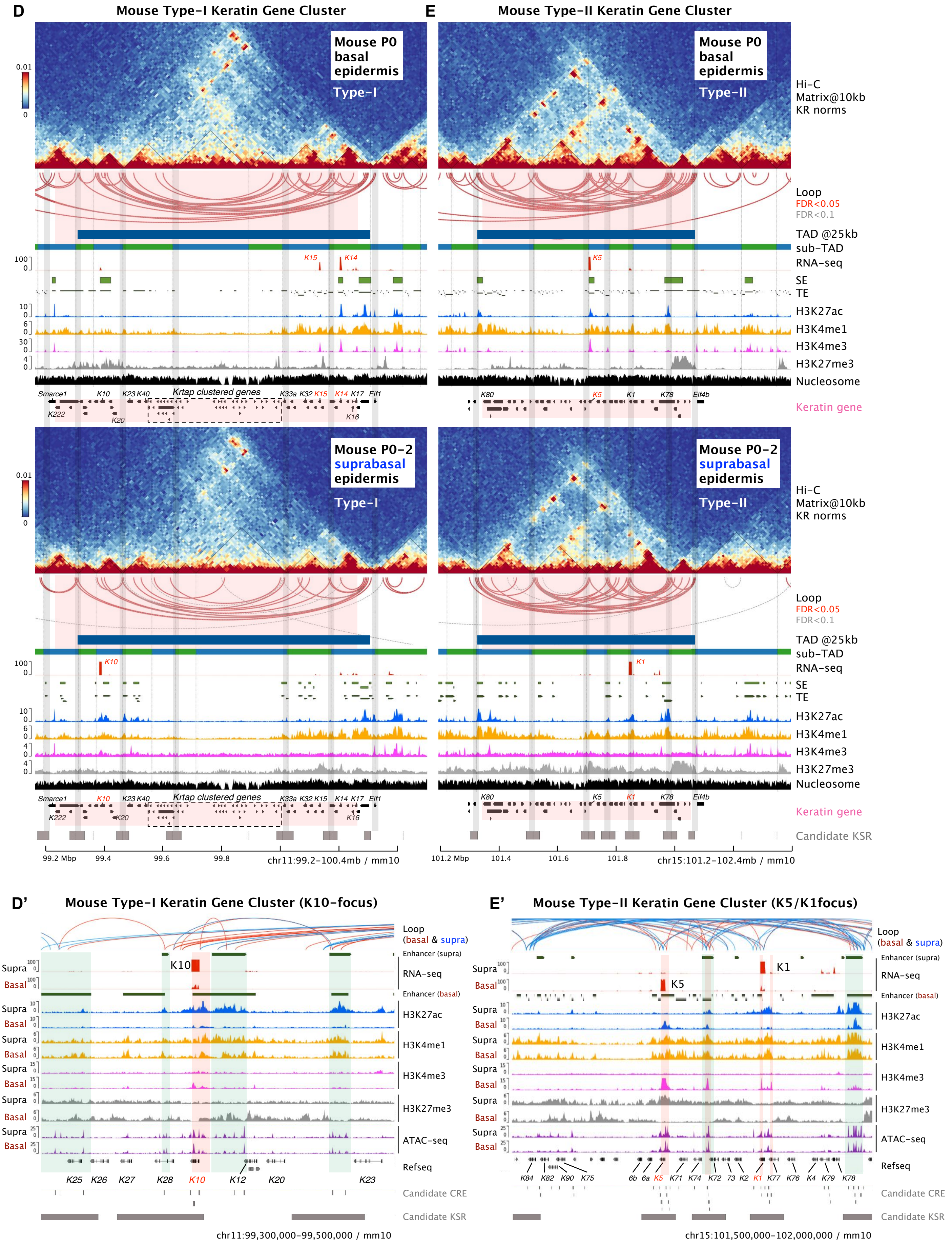

D
Mouse Type-I Keratin Gene Cluster
E
Mouse Type-II Keratin Gene Cluster
Mouse P0 basal epidermis
Type-I
Type-II
Hi-C Matrix@10kb KR norms
Loop
FDR<0.05
FDR<0.1
TAD @25kb
sub-TAD
RNA-seq
SE
TE
H3K27ac
H3K4me1
H3K4me3
H3K27me3
Nucleosome
Keratin gene
Smarce1
K10
K23 K40
Krtap clustered genes
K33a K32 K15
K14 K17 Eif1
K222
K20
K16
K80
K5
K1
K78
Eif4b
Mouse P0-2 suprabasal epidermis
Candidate KSR
99.2 Mbp
99.4
99.6
99.8
chr11:99.2-100.4mb / mm10
101.2 Mbp
101.4
101.6
101.8
chr15:101.2-102.4mb / mm10
D'
Mouse Type-I Keratin Gene Cluster (K10-focus)
E'
Mouse Type-II Keratin Gene Cluster (K5/K1focus)
Loop
(basal & supra)
Enhancer (supra)
Enhancer (basal)
Supra
Basal
ATAC-seq
Refseq
K25 K26 K27 K28 K10 K12 K20 K23
K84 K82 K90 K75 6b 6a K5 K71 K74 K72 73 K2 K1 K77 K76 K4 K79 K78
Candidate CRE
chr11:99,300,000-99,500,000 / mm10
chr15:101,500,000-102,000,000 / mm10

Figure S3 (F-G)

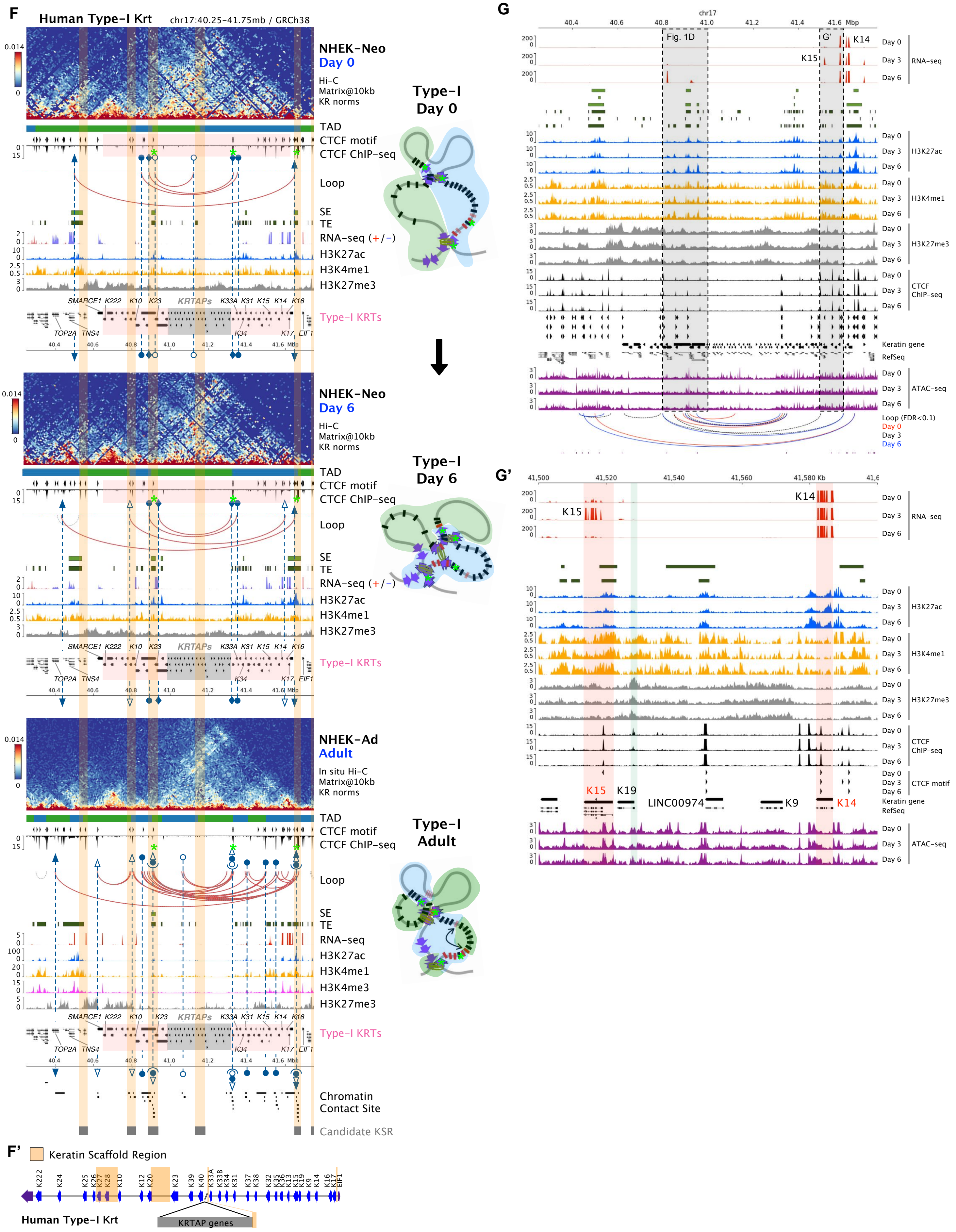
F
Human Type-I Krt
chr17:40.25-41.75mb / GRCh38
NHEK-Neo
Day 0
Hi-C
Matrix@10kb
KR norms
TAD
CTCF motif
CTCF ChIP-seq
Loop
SE
TE
RNA-seq (+/−)
H3K27ac
H3K4me1
H3K27me3
SMARCE1 K222 K10 K23 KRTAPs K33A K31 K15 K14 K16
TOP2A TNS4 K34 K17 EIF1
Type-I KRTs
Type-I
Day 0
NHEK-Neo
Day 6
Type-I
Day 6
NHEK-Ad
Adult
In situ Hi-C
Matrix@10kb
KR norms
H3K4me3
Type-I
Adult
Chromatin
Contact Site
Candidate KSR
F'
Keratin Scaffold Region
Human Type-I Krt
KRTAP genes
G
chr17
Fig. 1D
G'
K14
K15
Day 0
Day 3
Day 6
RNA-seq
H3K27ac
H3K4me1
H3K27me3
CTCF
ChIP-seq
Keratin gene
RefSeq
ATAC-seq
Loop (FDR<0.1)
G'
K15
K19
LINC00974
K9
K14
CTCF motif

Figure S3 (H-I)

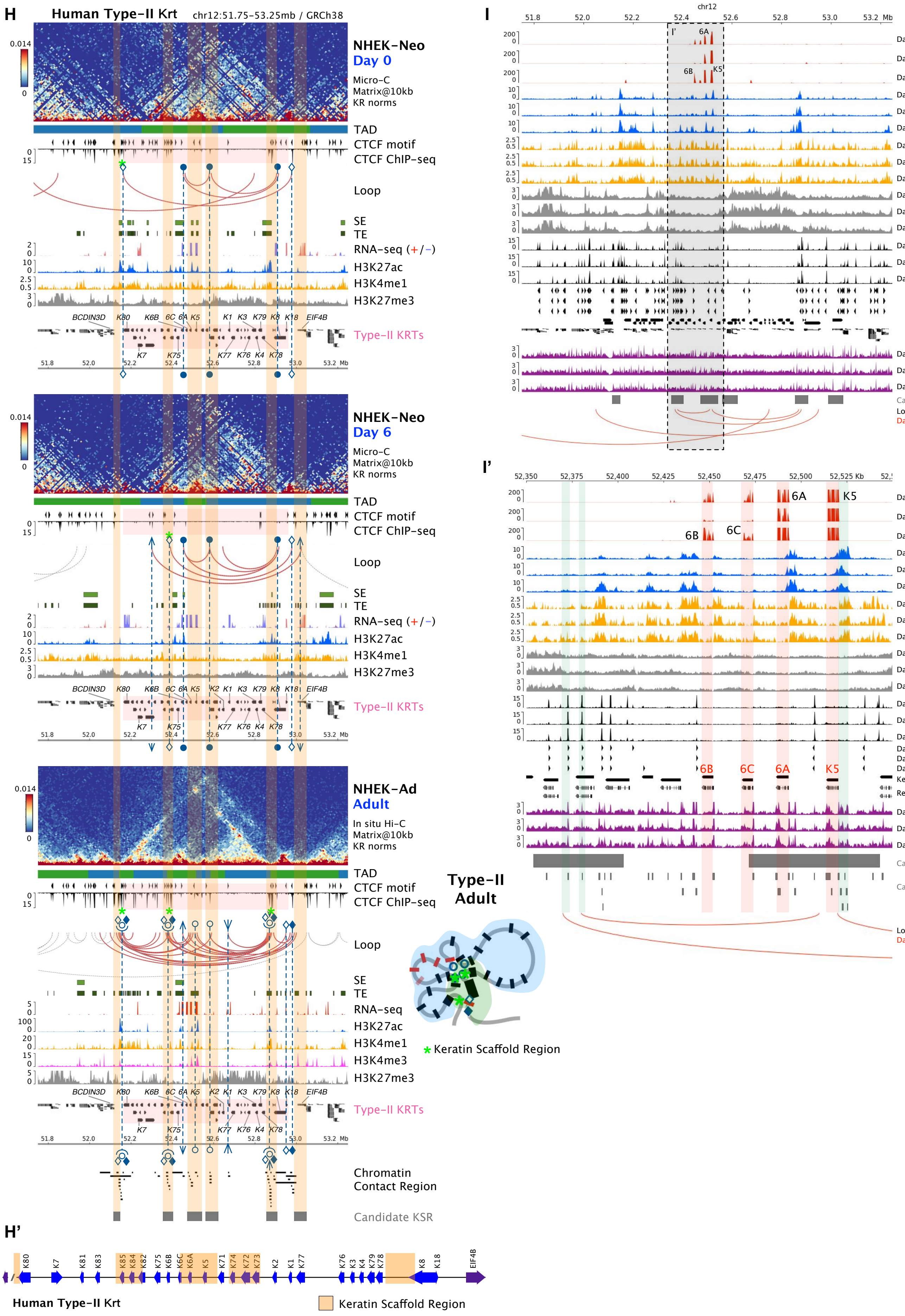
H
Human Type-II Krt
chr12:51.75-53.25mb / GRCh38
NHEK-Neo
Day 0
Micro-C
Matrix@10kb
KR norms
TAD
CTCF motif
CTCF ChIP-seq
Loop
SE
TE
RNA-seq (+/−)
H3K27ac
H3K4me1
H3K27me3
Type-II KRTs
NHEK-Neo
Day 6
NHEK-Ad
Adult
In situ Hi-C
Matrix@10kb
KR norms
H3K4me3
Chromatin
Contact Region
Candidate KSR
Type-II
Adult
Keratin Scaffold Region
H'
Human Type-II Krt
Keratin Scaffold Region
I
chr12
RNA-seq
H3K27ac
H3K4me1
H3K27me3
CTCF ChIP-seq
ATAC-seq
Candidate KSR
Loop (FDR<0.1)
Day 0
I'
CTCF motif
Keratin gene
RefSeq
Candidate CRE

Figure S3 (J-Q)

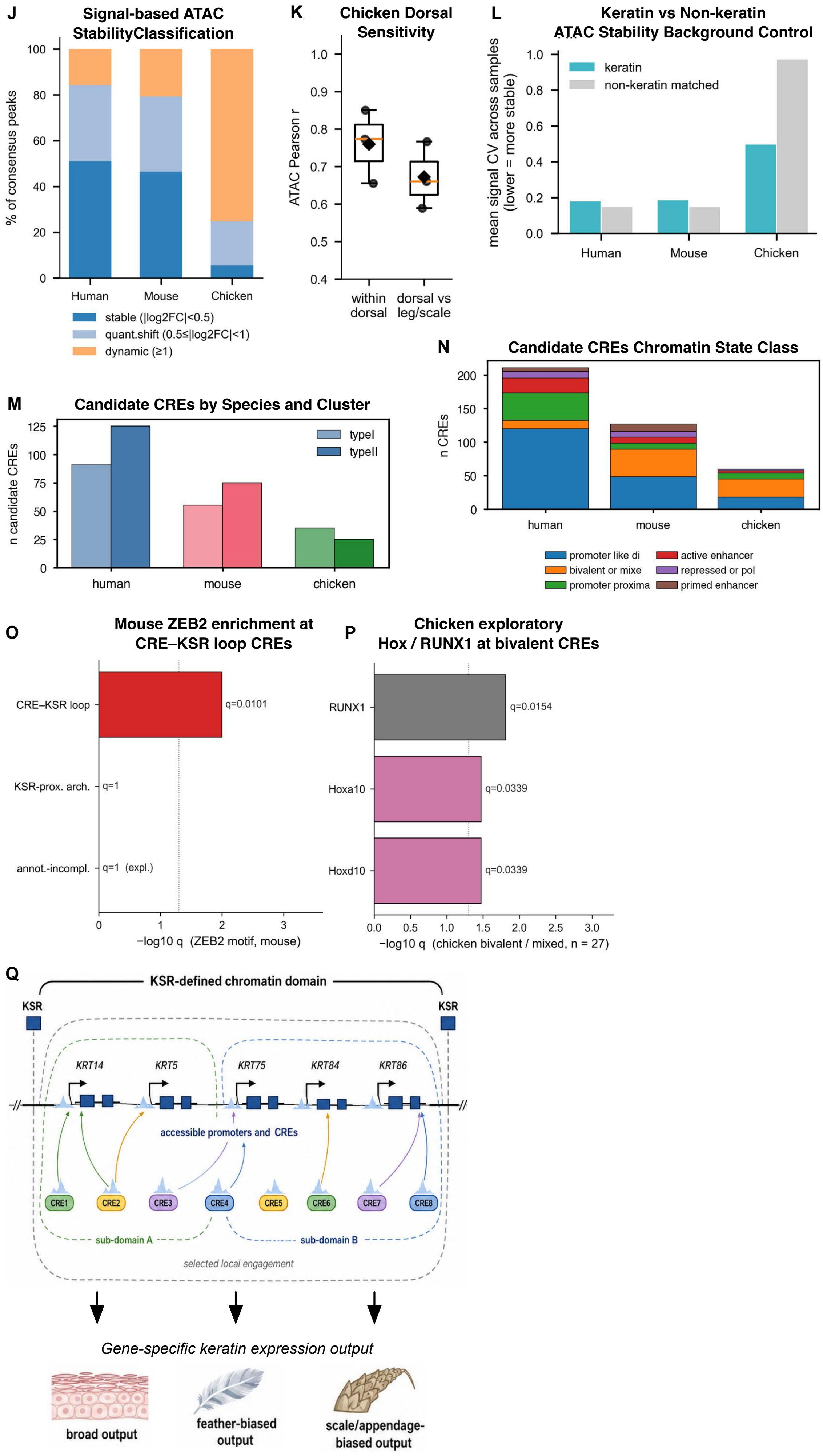
J
Signal-based ATAC StabilityClassification
% of consensus peaks
Human
Mouse
Chicken
stable (|log2FC|<0.5)
quant.shift (0.5≤|log2FC|<1)
dynamic (≥1)
K
Chicken Dorsal Sensitivity
ATAC Pearson r
within dorsal
dorsal vs leg/scale
L
Keratin vs Non-keratin ATAC Stability Background Control
mean signal CV across samples (lower = more stable)
keratin
non-keratin matched
Human
Mouse
Chicken
M
Candidate CREs by Species and Cluster
n candidate CREs
typeI
typeII
human
mouse
chicken
N
Candidate CREs Chromatin State Class
n CREs
human
mouse
chicken
promoter like di
bivalent or mixe
promoter proxima
active enhancer
repressed or pol
primed enhancer
O
Mouse ZEB2 enrichment at CRE–KSR loop CREs
CRE–KSR loop
q=0.0101
KSR-prox. arch.
q=1
annot.-incompl.
q=1 (expl.)
−log10 q (ZEB2 motif, mouse)
P
Chicken exploratory Hox / RUNX1 at bivalent CREs
RUNX1
q=0.0154
Hoxa10
q=0.0339
Hoxd10
q=0.0339
−log10 q (chicken bivalent / mixed, n = 27)
Q
KSR-defined chromatin domain
KSR
KSR
KRT14
KRT5
KRT75
KRT84
KRT86
accessible promoters and CREs
CRE1
CRE2
CRE3
CRE4
CRE5
CRE6
CRE7
CRE8
sub-domain A
sub-domain B
selected local engagement
Gene-specific keratin expression output
broad output
feather-biased output
scale/appendage-biased output

Figure S4

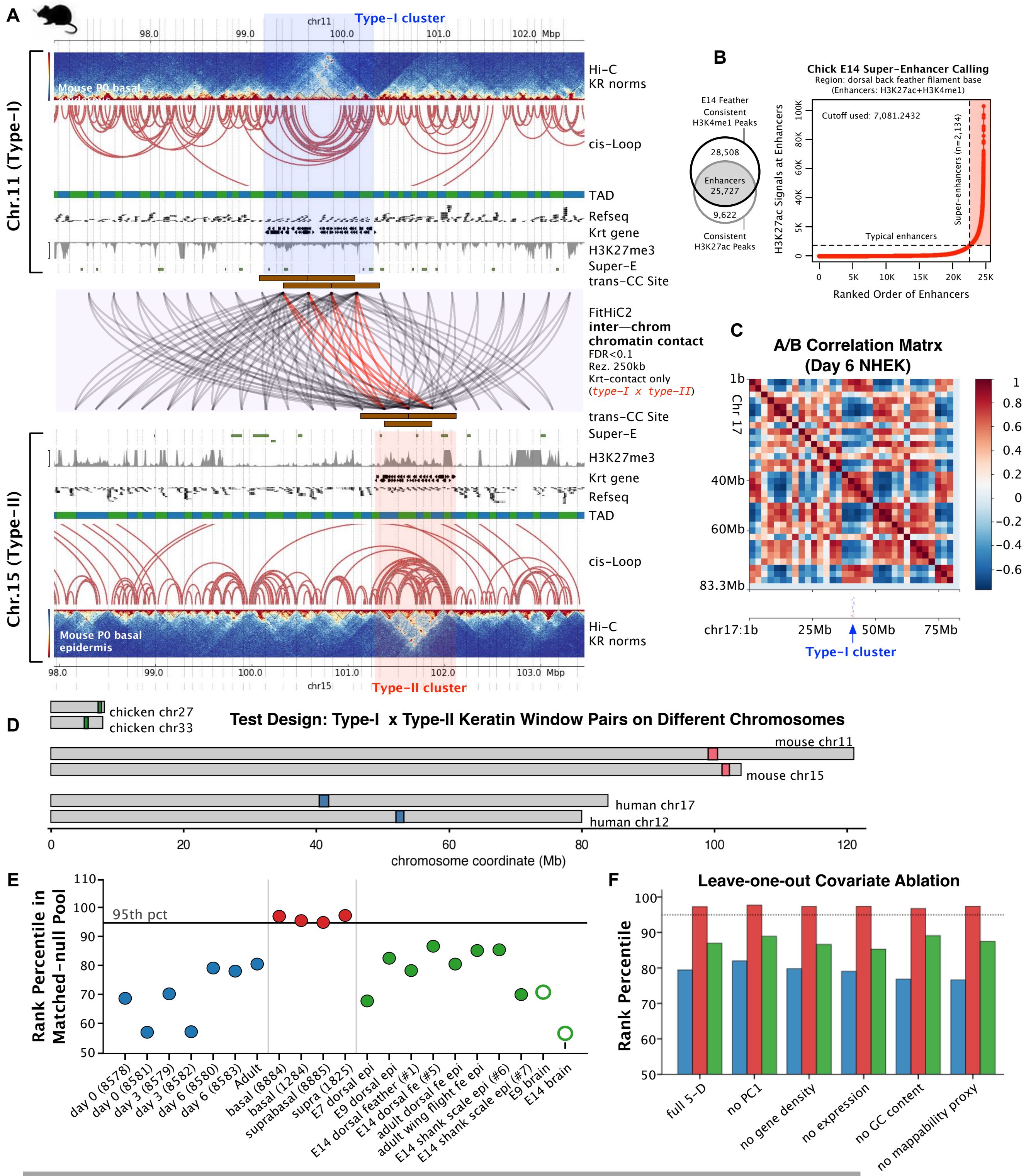


**G**

| | Clustered−Enhancer Name | ATAC Signal | MATCH−Predicted Transcription Factor Binding Sites (Factor genes have RNA−seq signals at exons and/or ATAC−seq signals around their core promoters) |
|---|---|---|---|
| Type−I α−Keratin Gene Cluster | **E14 feather Super−Enhancer (SE1)** chr27:7290775−7396392 | P1* | KLF6, ING4, BPTF, GTF2IRD1, NF1A, SMAD factors, ZSCAN4 (possible chicken orthologs: LOC107052489 and ZKSCAN7L), KLF4 |
| | | P2 | Sp100 (no chicken ortholog identified), KLF6, RNF96 (chicken ortholog: TRIM1BL), ZFP161, GTF2IRD1, YY1−like factors (chicken ortholog: YY2), E2f1 |
| | | P3* | KLF6, ING4, GTF2IRD1, TFCP2, ZFP161, KLF4, AhR/ARNT, NFAT−related factors, HSF dimer, CTCF, Mafa group |
| | | P4* | KLF6, GTF2IRD1, CTCF, RNF96 (chicken ortholog: TRIM1BL), Ets−related factors, MafA group, Pbx, KLF4, ZFP161 |
| | | P5* | CTCF, ZSCAN4 (possible chicken orthologs: LOC107052489 and ZKSCAN7L), HSF dimer, TEF, KLF6, GTF2IRD1, GR−like receptor (chicken: Nr3c1), p53−related factors, ZBTB14, NF1A, MAF group, MafA group, NFAT−related factors, HIC1 |
| Type−II α−Keratin Gene Cluster | **E14 feather Super−Enhancer (SE2)** chr33:5163545−5220612 | P6 | CP2−related factors, Ikaros, GATA, Sox−related factors, BPTF, GTF2IRD1, ING4, MAF group, TEF, AhR/ARNT, GR−like receptors, SREBP factors |
| | | P7 | KLF6, SMAD factors, Ikaros, AP−1, MafA group, REST, ING4, AhR/ARNT, RBP−Jkappa, ER group (e.g. PPARG and RAR, low/no Esr1/2), RREB−1, GLI group |
| | | P8 | GATA, Kaiso, KLF6, SREBP factors, AhR/ARNT, NFAT−related factors, KLF4 group, MAF group, RREB−1 |
